# *Arabidopsis* iron superoxide dismutase 1 protects against methyl viologen-induced oxidative stress in a copper-dependent manner

**DOI:** 10.1101/2021.09.20.461038

**Authors:** Pavol Melicher, Petr Dvořák, Yuliya Krasylenko, Alexey Shapiguzov, Jaakko Kangasjärvi, Jozef Šamaj, Tomáš Takáč

**Affiliations:** Department of Cell Biology, Centre of the Region Haná for Biotechnological and Agricultural Research, Faculty of Science, Palacký University Olomouc, Šlechtitelů 27, Olomouc, 783 71, Czech Republic; Organismal and Evolutionary Biology Research Programme, Faculty of Biological and Environmental Sciences, and Viikki Plant Science Center, University of Helsinki, FI-00014 Helsinki, Finland; Institute of Plant Physiology, Russian Academy of Sciences, Botanicheskaya Street, 35, 127276 Moscow, Russia; Natural Resources Institute Finland (Luke), Production Systems, Toivonlinnantie 518, FI-21500 Piikkiö, Finland

**Keywords:** superoxide dismutase, FSD1, *Arabidopsis*, methyl viologen, proteomics, copper, ferredoxin, protein carbonylation, oxidative stress

## Abstract

Iron superoxide dismutase 1 (FSD1) was recently characterized as a plastidial, cytoplasmic, and nuclear superoxide dismutase with osmoprotective and antioxidative functions. However, its role in oxidative stress tolerance is not well understood. Here, we characterized the role of FSD1 in response to methyl viologen (MV)-induced oxidative stress in *Arabidopsis thaliana*. The findings demonstrated that the antioxidative function of FSD1 depends on the availability of Cu^2+^ in growth media. Prolonged MV exposure led to a decreased accumulation rate of superoxide, higher levels of hydrogen peroxide production, and higher protein carbonylation in the *fsd1* mutants and transgenic plants lacking a plastidial pool of FSD1, compared to the wild type. MV led to a rapid increase in FSD1 activity, followed by a decrease. Chloroplastic localization of FSD1 is necessary for these changes. Proteomic analysis showed that the sensitivity of the *fsd1* mutants coincided with decreased abundance of ferredoxin and light PSII harvesting complex proteins, with altered levels of signaling proteins. Collectively, the study provides evidence for the conditional antioxidative function of FSD1 and its possible role in signaling.

## Introduction

Photosynthetic light reactions are inherently accompanied by the formation of reactive oxygen species (ROS) [1,2]. Superoxide anion radicals (O_2_^•–^) and singlet oxygen (^1^O_2_) are the primary ROS, while O_2_^•–^ may be converted to other ROS, such as hydrogen peroxide (H_2_ O_2_) or hydroxyl radicals (^•^OH) [3]. Under normal conditions, most chloroplastic O_2_^•–^ is produced on the acceptor site of photosystem I (PSI) [4,5]. Ferredoxin (Fd), the last electron acceptor of PSI, has been reported to produce O_2_^•–^; however, the reduction of O_2_ by Fd is highly dependent on available NADP^+^ and is lower than the contribution of O_2_ reduction by membrane-bound photosynthetic electron transport chain (PETC) components [6].

Methyl viologen (1,1’-dimethyl-4,4’-bipyridinium dichloride; MV), also known as paraquat, is a ROS-inducing, rapidly acting, non-selective herbicide. Its toxicity is exerted by catalyzing light-dependent electron transfer from PSI to O_2_, resulting in O_2_^•–^ production [7–9]. With a midpoint redox potential (Em) of −446 mV, the divalent MV cation (MV^2+^) easily accepts a single electron, which leads to the formation of a reduced cation radical (MV^+•^). The reaction most probably takes place in PSI; however, the specific site from which MV^2+^ accepts the electron is not clear. It is suggested that either Fd, PSI, or both might be the donors of electrons to MV^2+^ [7]. Shortly after MV^+•^ is produced, it is oxidized by O_2_ to form O_2_^•–^. In addition, MV^2+^ is regenerated and can be reduced to MV^+•^ again, thus repeating the entire O_2_^•–^ evolution cycle [10,11]. Within hours of application, the ROS produced by MV can overcome the protective mechanisms of the cell and lead to serious damage [7,8]. Owing to these characteristics, MV is widely used as an oxidative stress-inducing agent in plant science.

MV affects various plant processes. It can lead to rapid membrane damage due to lipid peroxidation and induce cell death [12,13]. Reportedly, MV-induced ROS significantly affect photosynthesis; for example, it leads to oxidation of the photosystem II (PSII) core protein D1 [14] or inhibition of D1 translation [15]. Moreover, H_2_O_2_ can damage the water-splitting complex of PSII [16] and reduce the maximal photochemical efficiency of PSII (F_v_/F_m_) [14,17–19].

Plants have evolved an efficient antioxidant defense against O_2_^•–^ [20]. Superoxide dismutase (SOD; EC 1.15.1.1) is an antioxidant metalloenzyme that represents the first line of defense against ROS [21–23]. SODs catalyze the dismutation of O_2_^•–^ to O_2_ and less reactive H_2_O_2_. Several SODs, such as FeSOD, MnSOD, and Cu/ZnSOD, are found in plants based on the metal cofactors in their active sites. Three genes encoding FeSOD (*FSD1, FSD2*, and *FSD3*), one encoding MnSOD (*MSD1*), and three encoding Cu/ZnSOD (*CSD1, CSD2*, and *CSD3*), [24,25] have been identified in the genome of *Arabidopsis*. They are compartmentalized in mitochondria (*MSD1*; [26]), peroxisomes (*CSD3*; [24]), nucleus (*FSD1*, [27]), cytoplasm (*CSD1, FSD1* [24,27,28]), and soluble chloroplast stroma (*FSD1*, [27]), or associated with chloroplast thylakoids (*CSD2, FSD2*, and *FSD3*; [24,28]). Thus, all *Arabidopsis* FSD isoforms are localized in the chloroplast, whereas FSD1 shows triple localization in the cytosol, chloroplast, and nucleus [27–29]. FSD2 and FSD3 proteins are tightly attached to the stromal side of thylakoid membranes and form heterocomplexes in the chloroplast nucleoids. They possibly prevent DNA damage caused by O_2_^•–^, as indicated by the physical interaction of FSD3 with the plastid envelope DNA-binding protein (PEND) [28]. Both *fsd2* and *fsd3* mutants show a higher accumulation rate of O_2_^•–^ under standard conditions [28] and high sensitivity to enhanced illumination [30], whereas *fsd1* shows a similar response and O_2_^•–^ levels to the wild type (WT) [28,30]. In contrast, another study showed that *fsd1* mutants exhibited hypersensitivity to MV-induced oxidative stress [27]. Similarly, heterologous overexpression of *Arabidopsis FSD1* in tobacco and maize led to increased tolerance against MV. In maize it also increased growth rates [31,32]. These genetic studies have questioned the role of *FSD1* in oxidative stress tolerance.

The main factors affecting FSD1 abundance and enzymatic activity are the availability of Fe^2+^ [33], Cu^2+^ [34,35], nitrogen [36], and sucrose [37]. *FSD1* expression and protein abundance increase with low Cu^2+^ availability, in parallel with the decline in *CSD1* and *CSD2* expression, when Cu^2+^ is redirected into housekeeping proteins and compounds such as plastocyanin and cytochrome-c oxidase [25,34,38]. In contrast, under Cu^2+^ sufficiency, CSD1 and CSD2 abundances rises, while FSD1 abundance drops to minimal levels [34,35]. The well-characterized transcription factor SQUAMOSA PROMOTER-BINDING PROTEIN-LIKE 7 (SPL7) is considered a major component in the regulation of Cu^2+^ deficiency responses that tightly controls the expression of the above-mentioned SODs in a Cu^2+^-dependent manner [34,35]. Though, the antioxidant roles of FSD1 under different Cu^2+^ availability conditions are not fully understood. Here, we aimed to adress these roles and gain insight into the mechanisms of FSD1-dependent responses of *Arabidopsis* to MV.

## Materials and Methods

### Plant material and growth conditions

Mature seeds of *Arabidopsis thaliana* ecotype Col-0 (WT), *fsd1-*1, and *fsd1-2* mutants as well as *fsd1-1* mutants expressing *proFSD1::FSD1:GFP* (FSD1-GFP) or *proFSD1::GFP::FSD1* (GFP-FSD1) [27] were surface-sterilized and placed on a half-strength Murashige and Skoog (MS) medium [39], prepared using MS basal salt mixture (Duchefa, cat n. M0221). For Walz imaging pulse-amplitude-modulation (PAM) measurements, a half-strength MS medium was prepared manually with increasing concentrations of Cu^2+^ by applying 0-3 µM CuSO_4_·5H_2_O. Seeds on plates were stratified at 4°C for 2 days to synchronize germination. Seedlings were grown at 21°C and 70% humidity under a 16/8 h (light/dark) photoperiod with a photosynthetic photon flux (PPF) of 120 μmol m^−2^s^−1^ in an environmental chamber (Weiss Technik, USA) provided by cool white fluorescent linear tube light sources (Philips MasterTL-D Reflex 36 W, light flow 3350 lm, light efficiency 93 lm·W^−1^) for a maximum of 14 days.

### Pulse□amplitude□modulation chlorophyll fluorescence imaging

Ten-day-old seedlings were horizontally grown on MS salt-based medium [39] plates with modified final concentrations of Cu^2+^. The plates with seedlings were loaded with 3 μM MV dissolved in liquid half-strength MS supplemented with 0.05% Tween 20, kept overnight in darkness, and on the next day were used for measurements of chlorophyll fluorescence kinetics using a PAM fluorometer according to a previously published protocol [40]. The analysis was performed in triplicate. Statistical evaluation of the obtained data was carried out by one-way ANOVA test with posthoc Tukey HSD test available at an online web statistical calculator (http://astatsa.com/OneWay_Anova_with_TukeyHSD/), which was also used for statistical evaluation of results obtained in all subsequent experiments if not mentioned otherwise. The graphical plots were prepared using Microsoft Excel and finalized using PowerPoint software.

### MV treatment for biochemical, histochemical, and proteomic analyses

At least six seedlings of each line, grown vertically for 14 days, were transferred from solid media to 6-well microtiter plates. Each well contained 7 ml liquid half-strength MS medium supplemented with 1 µM MV. For the mock control, distilled water was used instead of MV, and seedlings were incubated under the same conditions as before. The MV effect was potentiated by an enhanced illumination regime (180 μmol·m^−2^·s^−1^). The incubation time ranged from 30 min to 8 h. To ensure that circadian oscillations of *FSD1* expression had no impact on the results of analyses, all treatments were terminated at the same time in the day (6 p.m.). After incubation, the seedlings were gently wiped with filter papers to remove excess media. Then they were either stained for histochemistry studies or immediately frozen in liquid nitrogen and stored at -80°C.

### Protein extraction for biochemical analyses

The frozen plant material was crushed in liquid nitrogen to a fine powder using a mortar and pestle and transferred to Eppendorf tubes. The powder (100 mg) was resuspended and homogenized with 200 µl extraction buffer containing 50 mM sodium phosphate (pH 7.8), 10% glycerol, and 2 mM ascorbate. The tubes were placed on ice for 30 min and vortexed occasionally. Subsequently, the extract was centrifuged at 13 000 × g for 20 min at 4°C, and the protein concentration of the supernatant was measured according to the Bradford method [41]. The native protein extract was used for the in-gel detection of SOD and spectrophotometric measurement of ascorbate peroxidase (APX) activity. For immunoblot analysis, native extracts were supplemented with 4 times concentrated Laemmli SDS buffer (LB) at a 3:1 ratio (sample: LB) and 5% (v/v) β-mercaptoethanol. Samples were boiled for 5 min at 95°C.

### Immunoblot analysis and analysis of SOD isozymes

Denatured protein extracts were separated by SDS-PAGE on 10% TGX Stain-Free™ (Bio-Rad) gels prepared according to the manufacturer’s instructions using a Stain-Free FastCast Acrylamide Kit, 10% (Bio-Rad). For each sample, 20 µg protein was loaded onto a gel and separated at 180 V for 50 min. Proteins were then transferred to a polyvinylidene difluoride (PVDF) membrane (GE Healthcare, Little Chalfont, United Kingdom) in a wet tank unit (Bio-Rad) at a constant current of 230 mA for 1.5 h on ice. To ensure a successful transfer of the proteins, the membrane was visualized by ChemiDoc MP Imaging System (Bio-Rad) using the “Stain-Free Blot” protocol in Image Lab software (Bio-Rad). The membrane was blocked with 5% (w/v) Blotting-Grade Blocker (Bio-Rad) in Tris-buffered saline (TBS, 100 mM Tris-HCl; 150 mM NaCl; pH 7.6) overnight and subsequently incubated with anti-FeSOD1 (AS06 125, diluted 1:3000), anti-APX (AS08 368, diluted 1:5000), anti-Ferritin (AS10 674, diluted 1:3000), and anti-Ferredoxin 2 (AS20 4433, diluted 1:3000) primary antibodies (Agrisera, Vännäs, Sweden) in TBS-T (TBS supplemented with 0.1% (v/v) Tween-20) and 1% (w/v) Blotting-Grade Blocker (Bio-Rad) at 4°C overnight. The membrane was washed five times with TBS-T for 10 min. Afterward, it was incubated with horseradish peroxidase-conjugated goat anti-rabbit IgG secondary antibody (Santa Cruz Biotechnology, Santa Cruz, CA, USA), diluted 1:5000 in TBS-T containing 1% (w/v) BSA for 1.5 h at room temperature. After five washing steps in TBS-T, the membrane was incubated with Clarity Western ECL substrate (Bio-Rad) for 2 min, and chemiluminescence was detected using ChemiDoc MP Imaging System (Bio-Rad) using the “Chemi” protocol in Image Lab software (Bio-Rad).

For SOD isozyme detection, native protein extracts were separated on 10% native PAGE gel at a constant 20 mA/gel. The visualization of isozymes was performed as described previously [42]. The gel was imaged using Image Scanner III (GE Healthcare) and ChemiDoc MP Imaging System (Bio-Rad).

For both activity and immunoblotting analyses, band optical densities were measured using ImageJ software. All analyses were performed in triplicate.

### Spectrophotometric measurement of APX activity

To measure APX activity, the native protein extracts containing 10 µg protein were used. APX activity was analyzed as described previously [43] by measuring the H_2_O_2_-dependent oxidation of ascorbic acid at 290 nm. Absorbance was detected for 5 min at 10 s intervals. A molar extinction coefficient of 2.8 mM^−1^.cm^−1^ was used to calculate the change in the ascorbic acid content during the experiment.

### Histochemical detection of O_2_•− and H_2_O_2_

O _2_^•–^ and H_2_ O_2_ were visualized by histochemical staining using nitroblue tetrazolium (NBT) and diaminobenzidine (DAB), respectively [42]. DAB- and NBT-stained seedlings were transferred to a storage mixture containing 20% (v/v) glycerol and 80% (v/v) ethanol before loading on glass slides. Whole leaf rosettes were then transferred to a glass slide in a drop of 100% glycerol and covered with a coverslip. Imaging of the stained leaf rosettes was performed using Image Scanner III (GE Healthcare). Staining intensity was measured using ImageJ. For staining intensity evaluation, the first two true leaves were used. Thirty and twenty leaves were analysed for NBT and DAB staining analysis, respectively.

### Detection of carbonylated proteins

For analysis of carbonylated proteins, plant material was homogenized in liquid nitrogen and proteins were extracted in 50 mM HEPES (pH 7.5) containing 75 mM NaCl, 1 mM EGTA, 1 mM NaF, 10% glycerol, 50 mM DTT, cOmplete™ Protease Inhibitor Cocktail (Roche) and PhosSTOP™ (Sigma-Aldrich). To analyze protein oxidation through carbonylated amino acid residues, we used a commercial OxyBlot™ kit (cat n. S7150, Merck KGaA, Darmstadt, Germany). Sample preparation, processing, and immunoblotting were performed according to the manufacturer’s instructions. Signals were visualized by chemiluminescence using the ChemiDoc MP Imaging System (Bio-Rad). The analysis was performed in two biological replicates.

### Proteomic analysis

Plants (WT, *fsd1-1*, and *fsd1-2* mutants) treated with 1 μM MV for 8 h were homogenized in liquid nitrogen, and the proteins were extracted by phenol extraction [44]. Mock-and MV-treated samples were analyzed in triplicate. Proteins were digested by in-solution digestion using sequencing-grade modified trypsin [44]. Digested peptides were resuspended in 15 μL formic acid, and 5 μL of the suspension was used for the analysis.

The LC-ESI-MS/MS analyses were performed on a nanoflow HPLC system (Easy-nLC1200, Thermo Fisher Scientific) coupled to an Orbitrap Fusion Lumos mass spectrometer (Thermo Fisher Scientific, Bremen, Germany) equipped with a nano-electrospray ionization source. Peptides were first loaded onto a trapping column and subsequently separated on a 15 cm C18 column (75 μm × 15 cm, ReproSil-Pur 5 μm 200 Å C18-AQ, Dr. Maisch HPLC GmbH, Ammerbuch-Entringen, Germany). The mobile phase consisted of water with 0.1% formic acid (solvent A) and acetonitrile/water (80:20 (v/v)) with 0.1% formic acid (solvent B). Peptides were eluted with a linear 110 min gradient from 5% to 21% solvent B in 62 min and then to 36% solvent B in 110 min, followed by a wash stage with 100% solvent B. MS data were acquired automatically using Thermo Xcalibur 4.1. software (Thermo Fisher Scientific). An information-dependent acquisition method consisted of an Orbitrap MS survey scan with a mass range of 300–1750 m/z followed by HCD fragmentation for the most intense peptide ions.

Data files were searched for protein identification using Proteome Discoverer 2.3 software (Thermo Fisher Scientific) connected to an in-house server running the Mascot 2.7.0 software (Matrix Science). Data were searched against the SwissProt (version 2019_11) database using the *A. thaliana* taxonomy filter. The following parameters were used: static modifications: carbamidomethyl (C)*, variable modifications: oxidation (M), acetyl (protein N-term), peptide mass tolerance: ± 10 ppm, fragment mass tolerance: ± 0.02 Da, maximum missed cleavages: 2. Methionine oxidation is a common modification during sample processing and is generally included in the search parameters. For quantitation, a minimum of two files containing the same features was obtained, and the unique peptides were used. The precursor abundance was based on the intensity. All peptides were used for normalization. ANOVA (adjusted p ≤ 0.05) was used to filter statistically significant results, applied to proteins exhibiting a fold change ≥ of 1.5. Proteins identified by one peptide were excluded from the analysis. Proteins present in all three replicates corresponding to the control proteome and absent in all three replicates of the test proteome were considered unique for the control proteome and vice versa.

### Bioinformatic evaluation of differential proteomes

Proteins showing different abundances between samples were classified using Gene Ontology (GO) annotation analysis using OmicsBox software (BioBam Bioinformatics, Valencia, Spain). BLAST was performed against the *A. thaliana* NCBI database, permitting 1 BLAST hits. The following parameters were used for annotation: E value hit filter1.0E^-6^; annotation cutoff: 55; GO weight: 5, GO Slim. The protein interaction network was identified in STRING [45] database with a minimum required interaction score of 0.55. The iron-sulfur cluster binding proteins from amino acid sequences of the differential proteomes of WT, *fsd1-1*, and *fsd1-2* mutants were predicted in MetalPredator [46].

## Results

### FSD1 protects PSII from MV-induced inhibition in a Cu^2+^-dependent manner

To evaluate the impact of FSD1 on photosynthetic performance upon MV treatment, we monitored the time course of quantum efficiency of PSII (Fv/Fm) using chlorophyll fluorescence in intact WT plants, *fsd1* mutants and *fsd1-1* mutant carrying either FSD1-GFP or GFP-FSD1 fusion proteins. A previous study has shown that the N-terminal fusion of GFP to FSD1 disturbs the plastid-targeting motive, and the GFP-FSD1 line lacks the plastidic pool of FSD1 [27], whereas the FSD1-GFP expressing line contains FSD1 in abundance, similar to WT [27]. Moreover, the study has also demonstrated that the abundance of FSD1 in the WT and both complemented lines varies inversely with Cu^2+^ concentration in the growth media [27]. Therefore, in this study, we estimated the quantum efficiency of PSII at different Cu^2+^ levels to unravel the effects of FSD1 on photosynthesis.

PAM measurements indicated that the sensitivity of the *fsd1* mutant and the GFP-FSD1 line to MV depended on Cu^2+^ levels in the growth medium, with the highest rate of PSII inhibition observed upon Cu^2+^ deficiency. At the same time, in the WT and FSD1-GFP line, PSII inhibition was less pronounced and was largely independent of Cu^2+^ concentrations. After 4 h of exposure to light and at a Cu^2+^ concentration of 0.01 μM or lower, the difference between the sensitive and tolerant genotypes was statistically significant (Figure 1). The quantum efficiency of PSII in the GFP-FSD1 line was slightly higher than that of the mutants, but the difference between them was not significant. The difference in PSII inhibition between the sensitive and tolerant genotypes was alleviated under higher Cu^2+^ concentrations and became indistinguishable in 0.5 – 3 μM Cu^2+^ (Figure 1). These results suggest that plastidic FSD1 protects the photosynthetic apparatus in *Arabidopsis* against MV only under low Cu^2+^ conditions.

**Figure 1.**
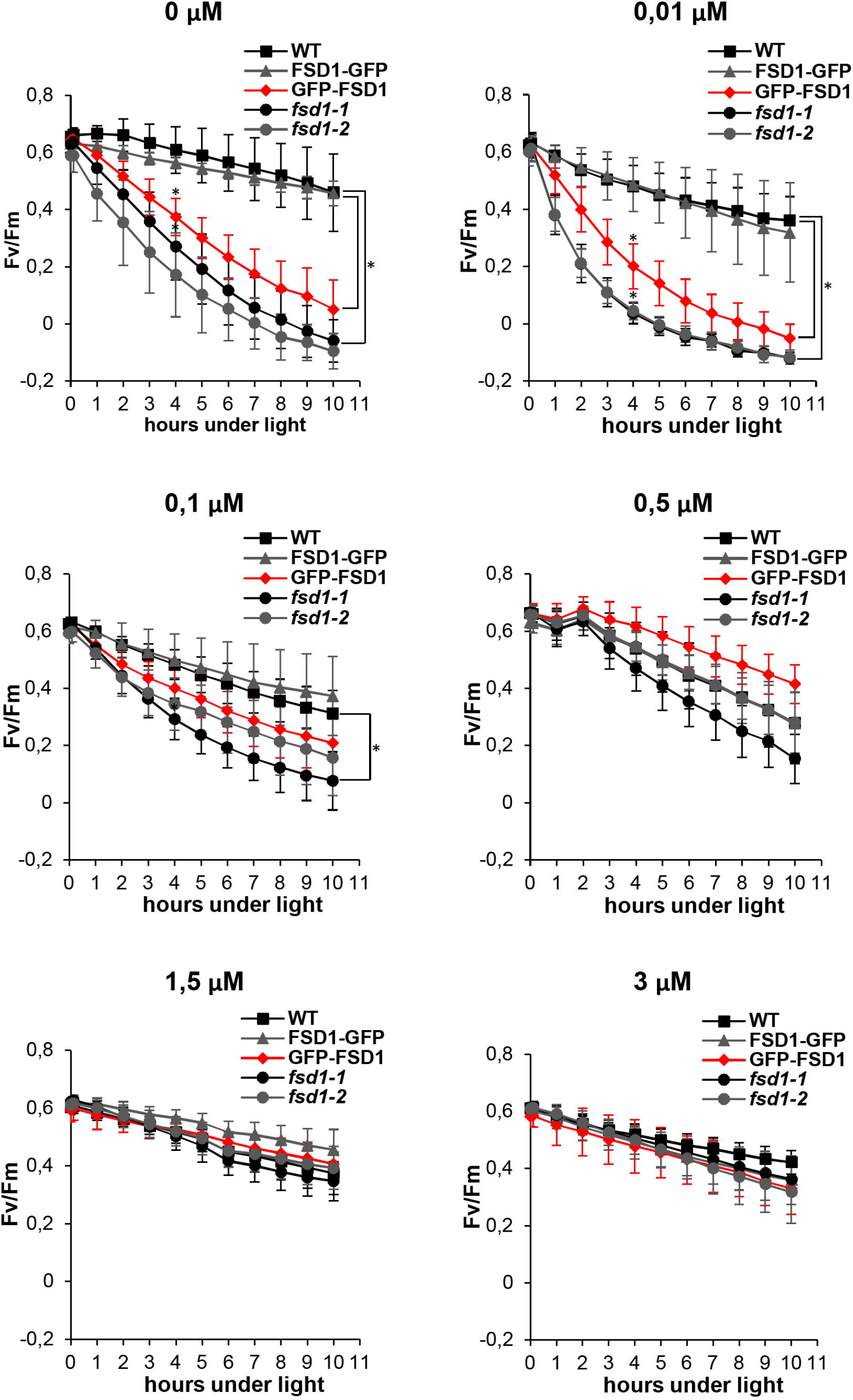
Effect of ROS production catalyzed by methyl viologen (MV) on the PSII activity (Fv/Fm) in wild type (WT), *fsd1* mutants and complemented *fsd1-1* mutants expressing FSD1-GFP or GFP-FSD1 in response to different concentrations of Cu^2+^ in the growing media. Each graph shows the time course of Fv/Fm values at different Cu^2+^ concentration, as indicated above each graph (mean ± SD, N = 9). Asterisks indicate a statistically significant difference between WT and *fsd1-1* or GFP-FSD1 lines as revealed by one-way ANOVA with post-hoc Tukey HSD test (p < 0.05).

### Chloroplast-localized FSD1 affects MV-induced ROS production

Histochemical staining of O_2_^•–^ using NBT (it forms a blue precipitate of formazan after reaction with O_2_^•–^) in WT, mutants, and transgenic lines subjected to MV treatment for 4 h demonstrated a decrease in formazan production, the extent of which differed among the studied lines (Figure 2A, S1A). In WT and FSD1-GFP lines, the staining intensity was slightly lower and accounted for more than 90% of the intensity detected in the mock control conditions. In *fsd1-1* and *fsd1-2* mutants, the decrease in staining intensity was much more pronounced and accounted for approximately 72% and 67% of the initial intensity, respectively. A similar change was observed in the GFP-FSD1 line (68% of the control). Therefore, a lower production rate of O_2_^•–^ was detected in MV-treated *fsd1* mutants and the GFP-FSD1 line than in WT and FSD1-GFP, highlighting the dependence of ROS metabolism on the presence of FSD1 in plastids.

**Figure 2.**
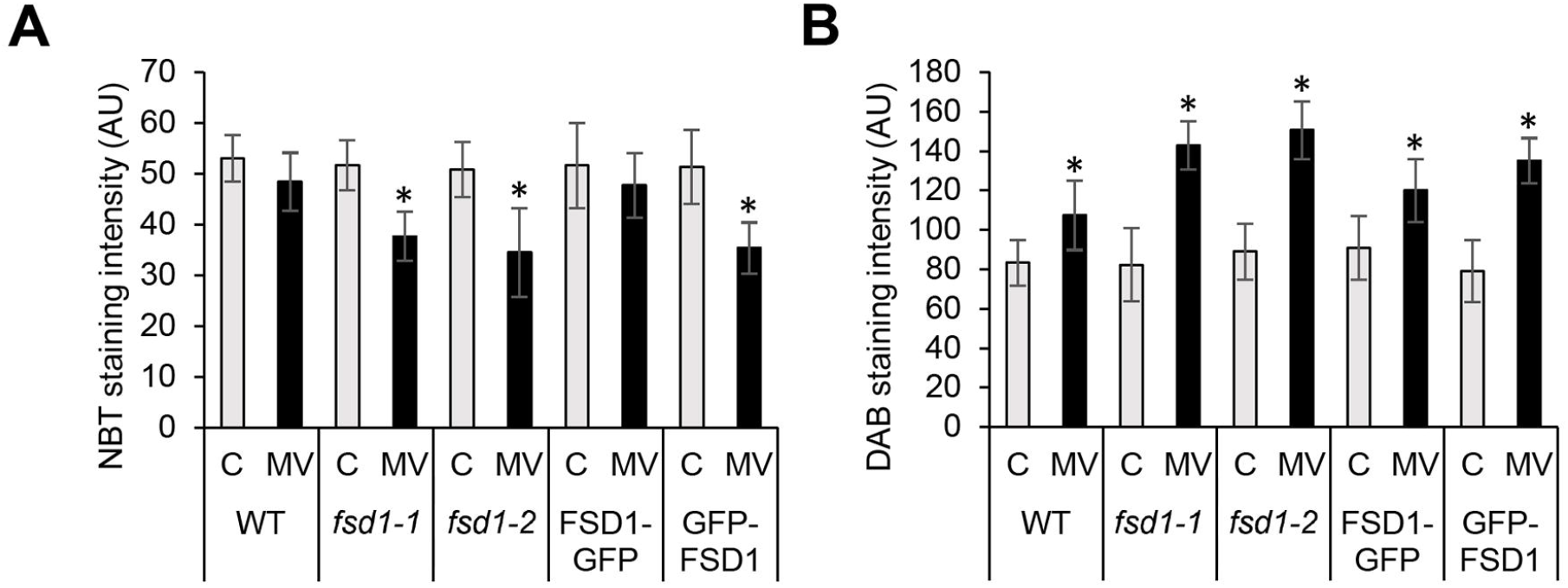
Semiquantitative analysis of superoxide (A) and hydrogen peroxide (B) staining in the first two true leaves of wild type plants (WT), *fsd1* mutants, FSD1-GFP and GFP-FSD1 lines in response to methyl viologen (MV) as estimated by quantification of signal intensity obtained by histochemical staining of leaves using nitroblue tetrazolium chloride (A) and 3,3′-diaminobenzidine (B). Asterisks indicate statistically significant difference at a p <0.05 as determined by one-way ANOVA with post-hoc Tukey HSD test (N = 30 for leaves stained by NBT and N = 20 for leaves stained by DAB).

Using DAB staining, we determined the production rate of H_2_O_2_ in the studied lines under MV treatment. The intensity of DAB staining was significantly increased in all the lines after 4 h of MV treatment depending on the presence of FSD1 in the plastids (Figure 2B, S1B). In the WT and FSD1-GFP line, the staining intensity was approximately 30% higher than that in the mock control; however, it was 70% higher in the *fsd1* mutants and the GFP-FSD1 line. These results strongly indicated the involvement of plastid-localized FSD1 in chloroplast ROS metabolism under MV treatment.

### Chloroplast-localized FSD1 affects the level of protein carbonylation under MV exposure

To monitor the rate of protein oxidation, we performed an oxyblot analysis that detected carbonyl groups by immunoblotting using a specific primary antibody. One hour of MV treatment did not lead to any significant changes in the intensity of the immunoreactive signal (Figure 3A), while the genotype-specific changes became apparent after 4 h (Figure 3B). In the WT, we observed only a slight increase of signal intensity. However, in the *fsd1* mutants and the GFP-FSD1 line, we observed a substantial increase, indicating a higher extent of protein oxidation. These results suggest that the sensitivity of plants to MV in terms of impact on the overall oxidative environment could be higher in the lines in which FSD1 is absent or not localized in plastids.

**Figure 3.**
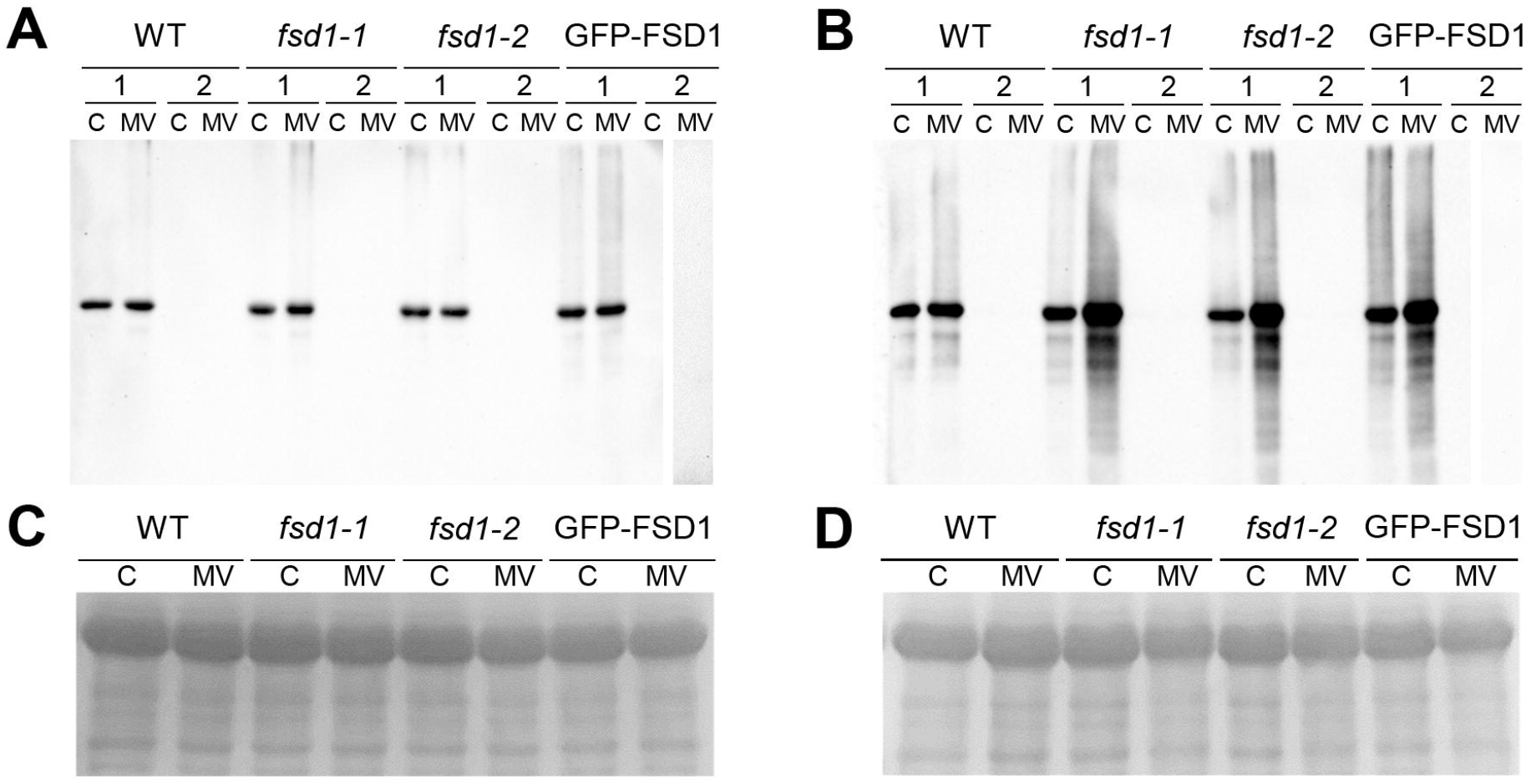
Detection of protein carbonylation in plants exposed to 1 µM methyl viologen (MV) and controls. (A, B) Detection of carbonyl groups in WT, *fsd1-1, fsd1-2* and GFP-FSD1 lines after 1 h (A) and 4 h (B) of MV treatment. Each blot contains protein extracts from mock (lane C) and 1 µM MV treated (lane MV) plants. Each sample was treated either with DNPH oxidation reagent (1) or a control reagent (2) included in the OxyBlot kit. The MV-treated samples of GFP-FSD1 derivatized with control reagent were run on separate gels followed by the same procedure. (C, D) Protein gels stained with Coomassie Brilliant Blue G-250 reagent demonstrating the equal amount of proteins loaded on gel for 1 h (C) and 4 h (D) treatment. Uncropped, full original images of the blots are documented in Figures S2 and S3.

### The response of FSD1 and APX to MV exposure

Next, we tested the response of FSD1 to MV treatment. The staining of SOD activity on native polyacrylamide gels allows differentiation among FSD1, MSD1, and Cu/ZnSOD activities [47]. Low copper levels in the media led to high FSD1 and low Cu/ZnSOD activities in the examined plants. These findings suggested that the specific response of FSD1 to MV can be determined while minimizing possible redundancy with CSD2 that is localized in plastids as well.

We observed that in WT, FSD1 abundance increased slightly after 30 min of MV treatment and decreased after 4 h (Figure 4A, B). However, FSD1 activity showed a more pronounced increase after 30 min and remained roughly constant after 4 h (Figure 4C, D). Similar results for FSD1 activity and abundance were observed in the FSD1-GFP line (data not shown). Strikingly, FSD1 activity and abundance were altered differently in the GFP-FSD1 line. Although the abundance of GFP-FSD1 did not substantially change throughout the experiment (Figure 4E, F), its activity remained unaffected after 30 min and 1 h of MV treatment, which was significantly inhibited after 4 h of treatment (Figure 4G, H). Taken together, these observations indicate that FSD1 activity and abundance are sensitive to MV and that its chloroplast localization is required for this sensitivity.

**Figure 4.**
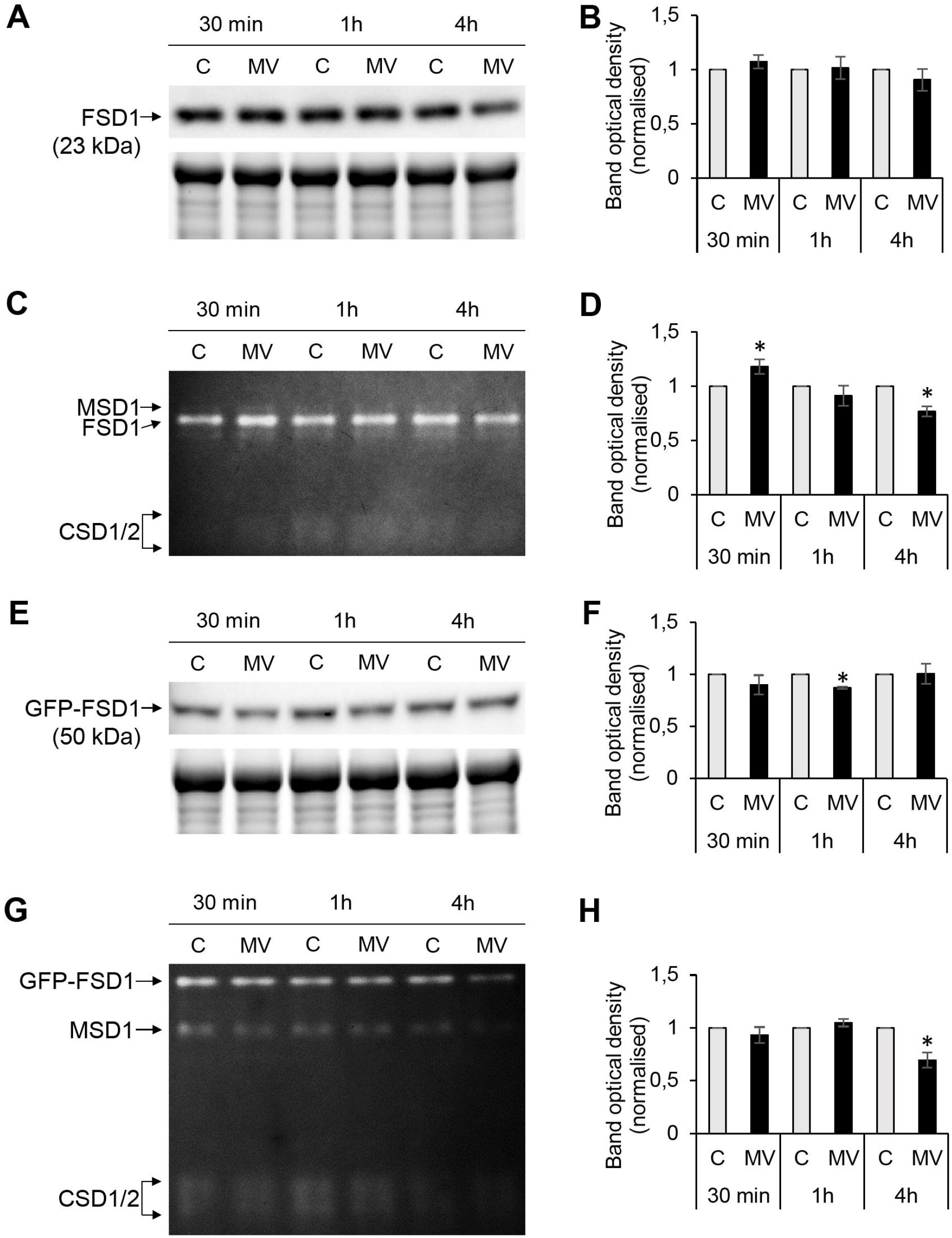
FSD1 abundance and activity in wild type (WT) (A-D) and GFP-FSD1 line (E-H) in response to methyl viologen (MV). (A, E) Immunoblots of FSD1 in WT (A) and GFP-FSD1 fusion protein in GFP-FSD1 line (E), supplemented with respective controls of protein loading using Stain-free gels. Each immunoblot contains protein extracts from mock (lane C) and 1 µM MV treated (lane MV) plants. (B, F) Quantification of band optical density in A and E, respectively. Values are expressed as relative to the mock control in each time point (mean ± SD, N = 3). (C, G) SOD activity staining in WT (C) and GFP-FSD1 line (G). Each gel contains protein extracts from mock (lane C) and 1 µM MV-treated (lane MV) plants. (D, H) Quantification of band optical density in C and G, respectively. Values are expressed as relative to the activity of FSD1 in mock control in each time point (mean ± SD, N = 3). Asterisks indicate a statistically significant difference between mock control and 1 µM MV treatment in designated time points as calculated by one-way ANOVA with post-hoc Tukey HSD test (asterisk indicates statistical significance at p < 0.05). Uncropped, full original images of the blots and gels are documented in Figures S4-S5.

APX is an enzyme with chloroplastic (thylakoids, stroma), cytoplasmic, and peroxisomal localizations and is responsible for the degradation of H_2_O_2_ [48]. Since ROS staining assays revealed altered H_2_O_2_ production rate in the studied lines (Figure 2), we next analyzed the APX activity and the abundance of cytosolic APX (cAPX) in these lines by spectrophotometric and immunoblotting approaches, respectively. The APX-specific activity in the mock control was similar among the studied lines (Figure 5A). After 30 min of MV treatment, APX activity increased by 21% in WT, while in *fsd1* mutants and GFP-FSD1 line, its activity remained unchanged (*fsd1-2*) or slightly decreased (*fsd1-1* and GFP-FSD1 line) compared to the control. Interestingly, there was no significant difference in the APX activity after 1 h among the studied lines (Figure 5A). In contrast, after 4 h of MV treatment, all lines showed a substantial decrease in APX activity (Figure 5A). These results indicate that the sensitivity of APX activity depends on the presence of FSD1 and its localization to plastids. In WT, *fsd1-1, fsd1-2* mutants, and GFP-FSD1 line, APX activity was decreased by 18%, 30%, 37%, and 33%, respectively, compared to that in the mock control (Figure 5A).

**Figure 5.**
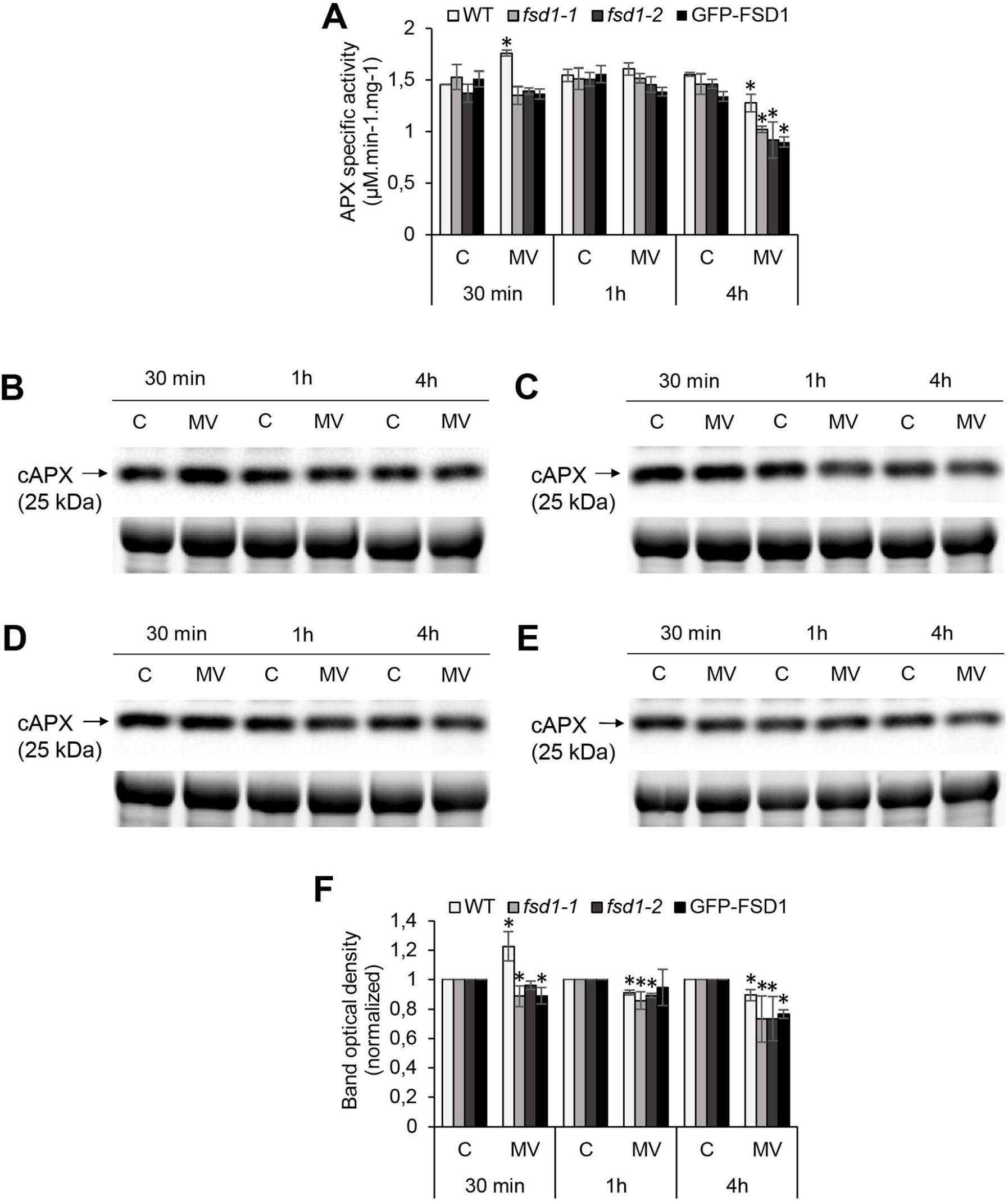
Analysis of ascorbate peroxidase (APX) activity and cytosolic APX (cAPX) abundance in wild type (WT), *fsd1-1 fsd1-2* and GFP-FSD1 lines after methyl viologen (MV) treatment. (A) Specific activity of APX measured spectrophotometrically. (B-E) Immunoblots of cAPX in wild type (WT; B), *fsd1-1* (C), *fsd1-2* (D) and GFP-FSD1 line (E), supplemented with respective controls of protein loading using Stain-free gels. Each immunoblot contains protein extracts from mock (lane C) and 1 µM MV treated (lane MV) plants. (F) Quantification of band optical density in B-E. Values are expressed as relative to the mock control in each time point (mean ± SD, N = 3). Asterisks indicate a statistically significant difference between mock and 1 µM MV treated plants in designated time points as calculated by one-way ANOVA with post-hoc Tukey HSD test (asterisk indicates statistical significance at p < 0.05). Uncropped, full original images of the blots and gels are documented in Figures S6-S7.

Anti-APX primary antibody allows the detection of cAPX by immunoblotting in our conditions. It recognizes two cAPX isoforms (APX1 and APX2) with a 0.5 kDa difference in molecular weight. Treatment with MV for 30 min caused a significant increase in cAPX abundance in WT, followed by a slight decrease after 1 h and a substantial decrease after 4 h (Figure 5B, C). In contrast, in the *fsd1* mutant lines, a slight decrease in abundance was observed as early as 30 min, followed by a more pronounced decrease after 1 h and 4 h (Figure 5B, D, E). Although the decrease in abundance after 4 h was higher in *fsd1* mutants than in WT, the difference between these lines was not significant. Similar to *fsd1* mutants, the effect of MV treatment on abundance decrease was also observed in the GFP-FSD1 line showing dependence on the FSD1 plastidial localization. However, the change in abundance after 1 h was negligible (Figure 5B, F).

Taken together, MV induced a fast transient increase in FSD1 and APX activity, as well as FSD1 and cAPX abundance. Notably, only plastidial FSD1 seemed responsive to short-term MV treatment. Prolonged MV exposure led to decreased abundances and activities of both FSD1 and APX.

### Proteomic analysis of fsd1 mutants after MV treatment

Next, we performed a comparative proteomic analysis of WT, *fsd1-1*, and *fsd1-2* lines exposed to 1 μM MV for 8 h to better understand the mechanisms underlying the increased sensitivity of *fsd1* mutants to MV. In addition to complete mass spectrometry/proteomics data deposited to PRIDE, the information pertinent to protein identification and quantification can be found in the supplemental material in the form of common Excel files given for each individual sample (Tables S1-S3). For each of the examined lines, we compared the proteomes of MV*-* and mock*-*treated plants. This analysis identified 65, 65, and 78 differentially abundant proteins in WT, *fsd1-1*, and *fsd1-2* plants, respectively (Figure 6A, Tables S1-S3). Of these, 29 proteins were commonly differentially regulated in both mutants, while 12 and 10 proteins were common between WT and *fsd1-1*, or WT and *fsd1-2*, respectively (Figure 6B), demonstrating the similarity between the two mutants.

**Figure 6.**
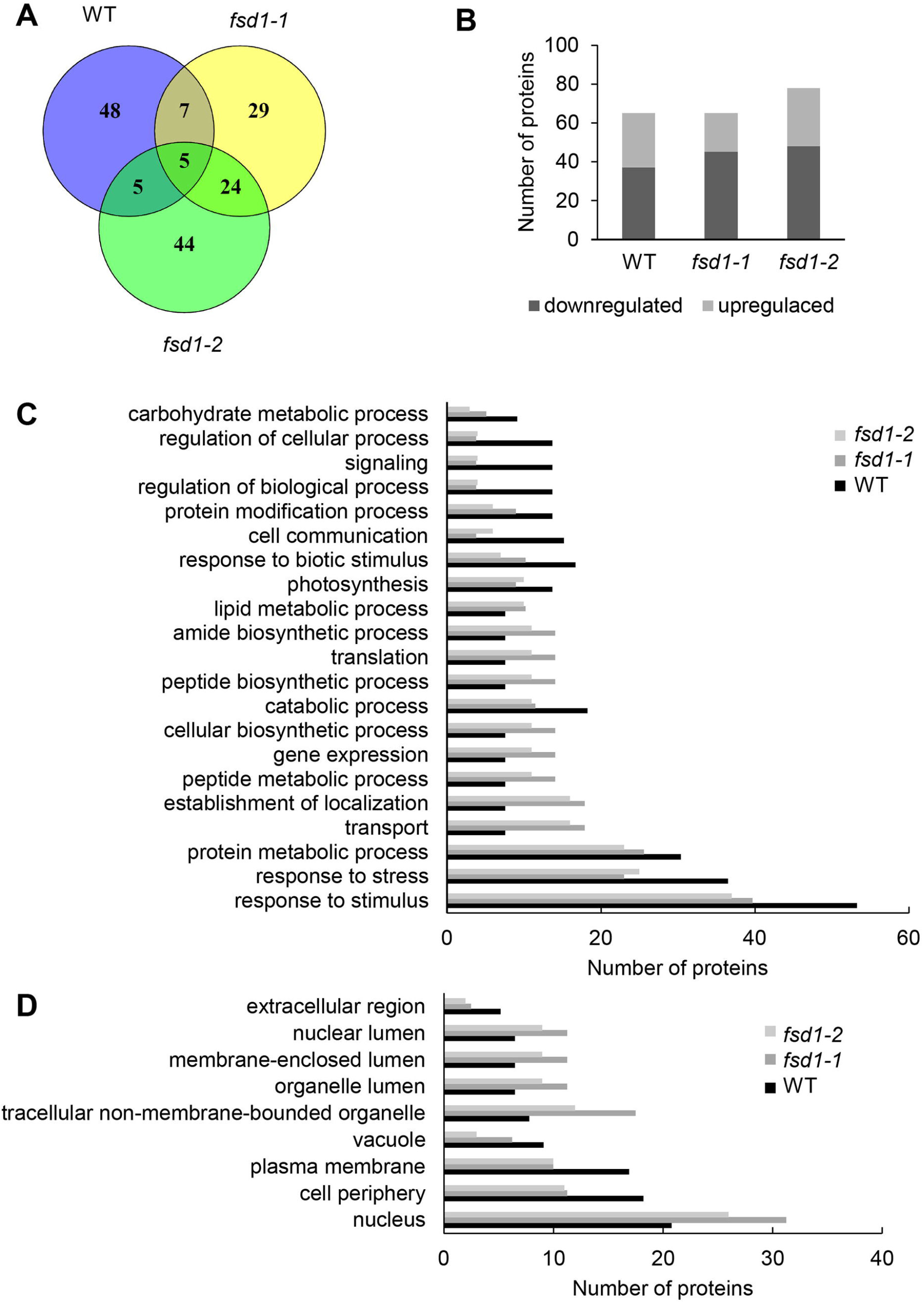
Evaluation of differentially abundant proteins (DAP) found in wild type (WT), *fsd1-1* and *fsd1-2* mutants exposed to 1 µM methyl viologen (MV) for 8 h. (A) Venn diagram showing the numbers of DAP. (B). Graph showing the comparison of numbers of upregulated and downregulated DAP. (C, D) Graph showing the abundance of gene ontology annotations found in the differential proteomes of the examined lines according to biological process (C), and cell compartment (D). Note, that in C and D, only those annotations, which differ among the lines are included.

First, we functionally categorized the differentially abundant proteins using GO annotations. Most changes were detected in GO categories for metabolism (i.e., primary, nitrogen compound, and protein metabolism) as well as the responses to abiotic stimuli and chemicals (Figure 6C). A smaller but significant change was seen in the following GO categories: responses to biotic, endogenous, and light stimuli, cell communication, signaling, gene expression, photosynthesis, and protein modification process (Figures 6C, S8). While some categories such as stress response, catabolism, photosynthesis, response to biotic stimuli, cell communication, and signaling were more abundant in the WT than in the mutants, other categories such as transport, peptide metabolic process, gene expression, and translation were more abundant in the mutants (Figure 6C). These findings suggest that FSD1 deficiency in the mutants limits the change in the abundance of proteins involved in stress response, signaling, photosynthesis, and proteins mediating cell*-*environment communication. Interestingly, MV led to the enrichment of proteins involved in biotic stress responses in the WT, but to a smaller extent in the mutants.

GO annotation according to the cell compartment showed that MV affected mainly cytoplasmic proteins and proteins localized in the plasma membrane and membranes of chloroplasts (thylakoids), nuclei, mitochondria, and the endomembrane system. Annotations such as nuclei and intracellular non-membrane-bounded organelles were enriched in mutants, while those named plasma membrane, vacuole, and extracellular region were more abundant in WT (Figures 6D, S9).

To gain insight into the possible mechanisms of FSD1-dependent MV tolerance, we focused on proteins involved in light reactions of photosynthesis (Table 1). WT plants showed an increased abundance of components of PSI, including the antenna protein LHCA3 and the plastocyanin major isoform. A similar increase in PSI component (PSI reaction center subunit VI-1 and subunit psaK) abundance was observed in *fsd1-2* mutants, suggesting that the abundance of these proteins was similarly affected in WT and the mutants. Substantial differences between the WT and mutants were observed in the abundance of Fd isoforms. While WT showed decreased levels of only one Fd isoform (C1), the mutants exhibited decreased levels of the four Fd isoforms (Table 1). The decreased abundance of Fd2 was validated by immunoblotting with an anti-Fd2 antibody (Figure 7A).

**Table 1.**
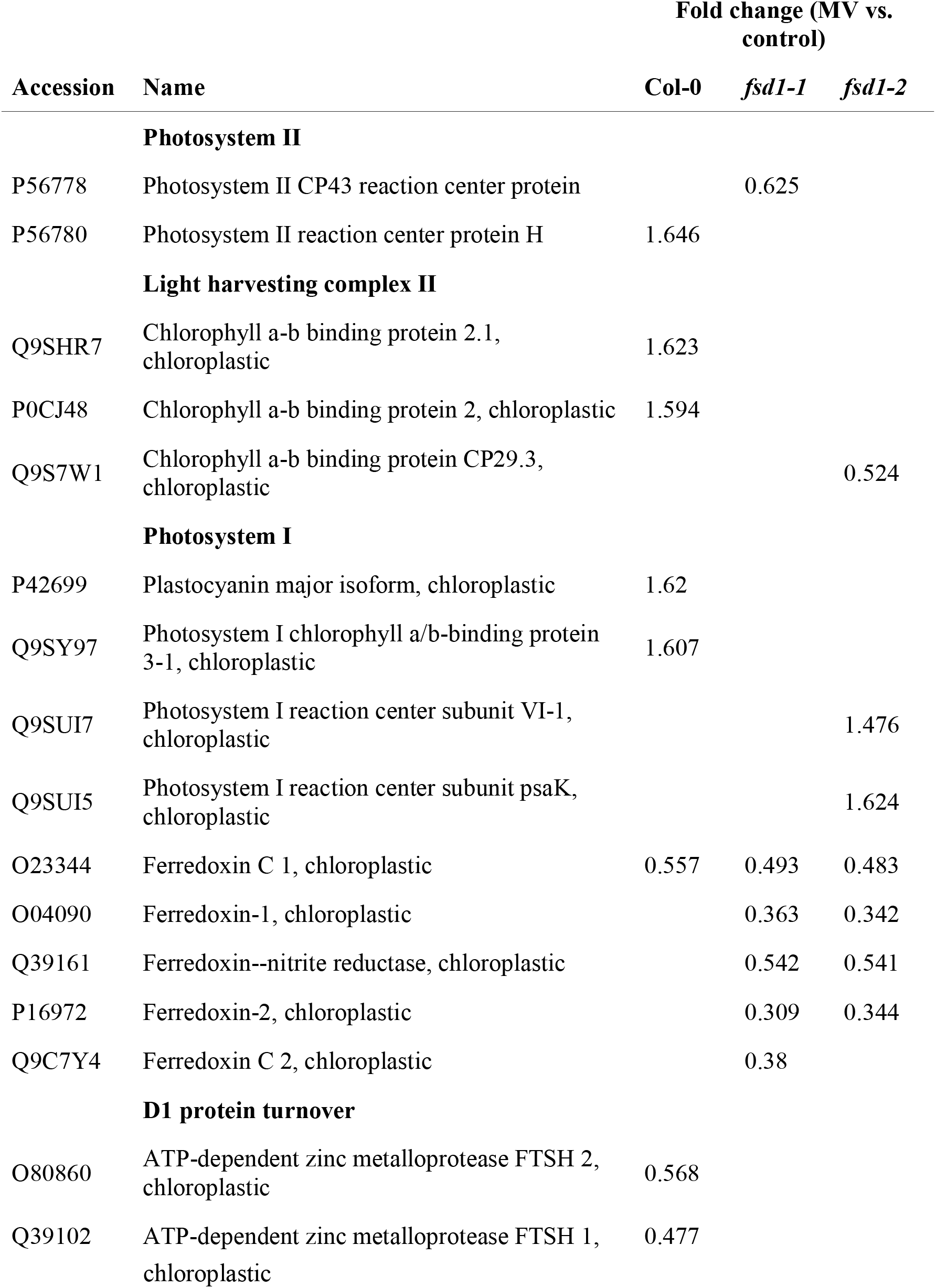
Proteins involved in light reactions of photosynthesis. Note that the empty cell means that the protein was not found in the differential proteome of a respective line.

**Figure 7.**
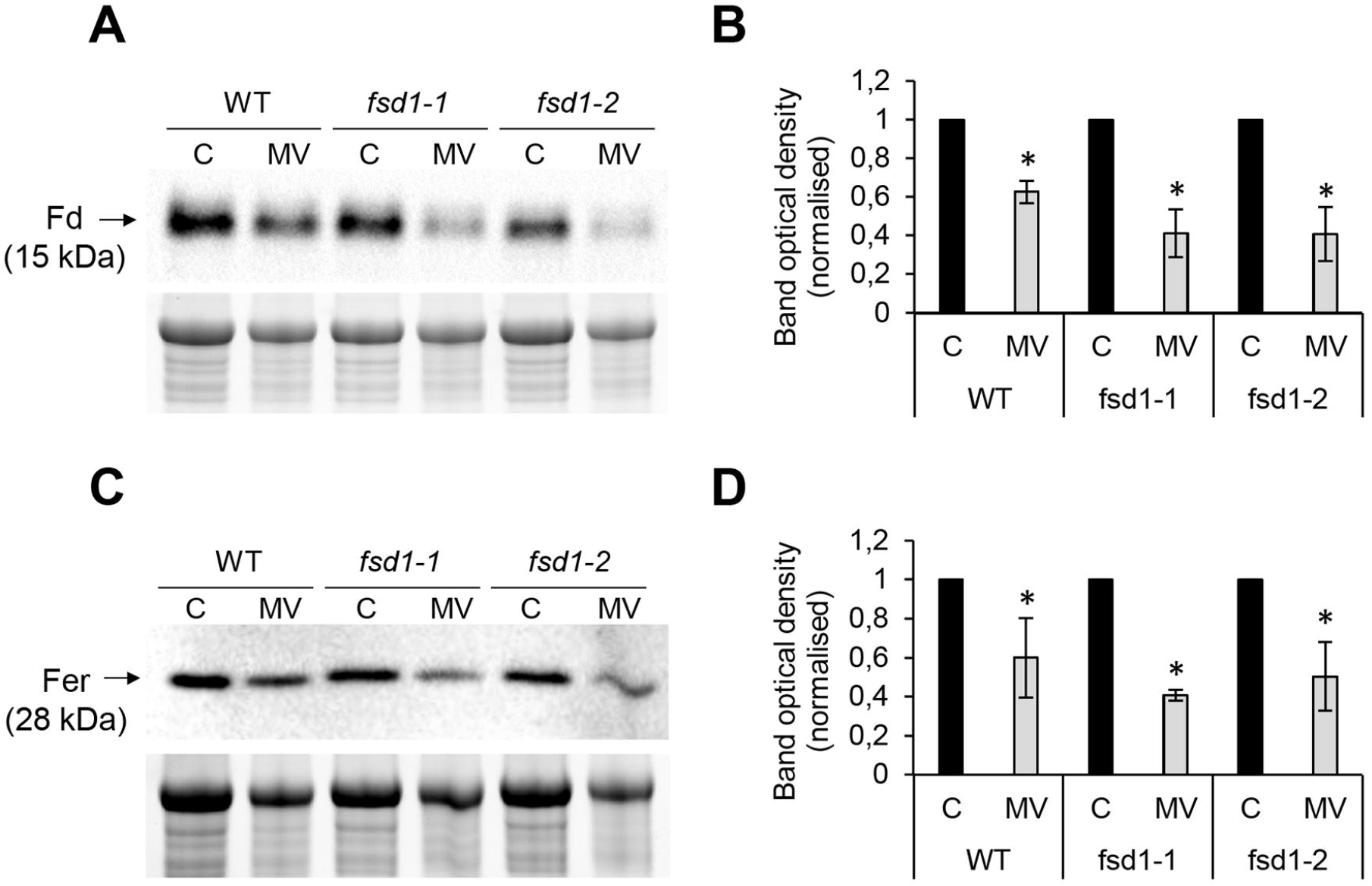
Immunoblotting analysis of ferredoxin 2 (Fd) and ferritins (Fer) abundance in wild type (WT) plants, *fsd1-1* and *fsd1-2* mutants in response to 8 h long methyl viologen (MV) treatment. Immunoblots of Fd (A) and Fer (C) in WT, *fsd1-1* and *fsd1-2*, supplemented with respective controls of protein loading using Stain-free gels. (B, D) Quantification of band optical densities in A and C. Values are expressed as relative to the mock control in each line (mean ± SD, N = 3). Each gel contains protein extracts from mock (lane C) and 1 µM MV-treated (lane MV) plants. Asterisks indicate a statistically significant difference between mock control and 1 µM MV treatment in 8 h as revealed by one-way ANOVA with post-hoc Tukey HSD test (p < 0.05). Uncropped, full original images of the blots are documented in Figures S13.

Another protein with decreased abundance in the mutants was Fd-nitrite reductase (Table 1). Isoforms of the chlorophyll *a-b* binding proteins, comprising the light-harvesting complex of PSII [49] were more abundant in WT and less abundant in the *fsd1-2* mutant. Moreover, ATP-dependent zinc metalloproteases, FTSH 1 and 2, were less abundant in WT, whereas their abundance did not differ among the mutants.

Fds contain Fe-S clusters as prosthetic groups in their structure. Since Fe-S clusters are known targets of O_2_^•–^ [50], we used MetalPredator software to predict the presence of proteins that bind Fe-S clusters in the differential proteomes (Table 2). Six, ten, and eight Fe-S cluster-binding proteins were affected in the WT, *fsd1-1*, and *fsd1-2* mutants, respectively. The majority of the predicted proteins showed a decreased abundance in response to MV. As noted above, mutants showed reduced levels of multiple Fd isoforms. Some other proteins, such as 3-isopropylmalate dehydratase large subunit and amidophosphoribosyltransferase 2, were equally under-represented in all lines. Notably, the CDGSH Fe-S domain-containing protein NEET showed a mutant-specific increase in abundance. These results demonstrate the overall increased sensitivity of Fe-S cluster-binding proteins in *fsd1* mutants.

**Table 2.**
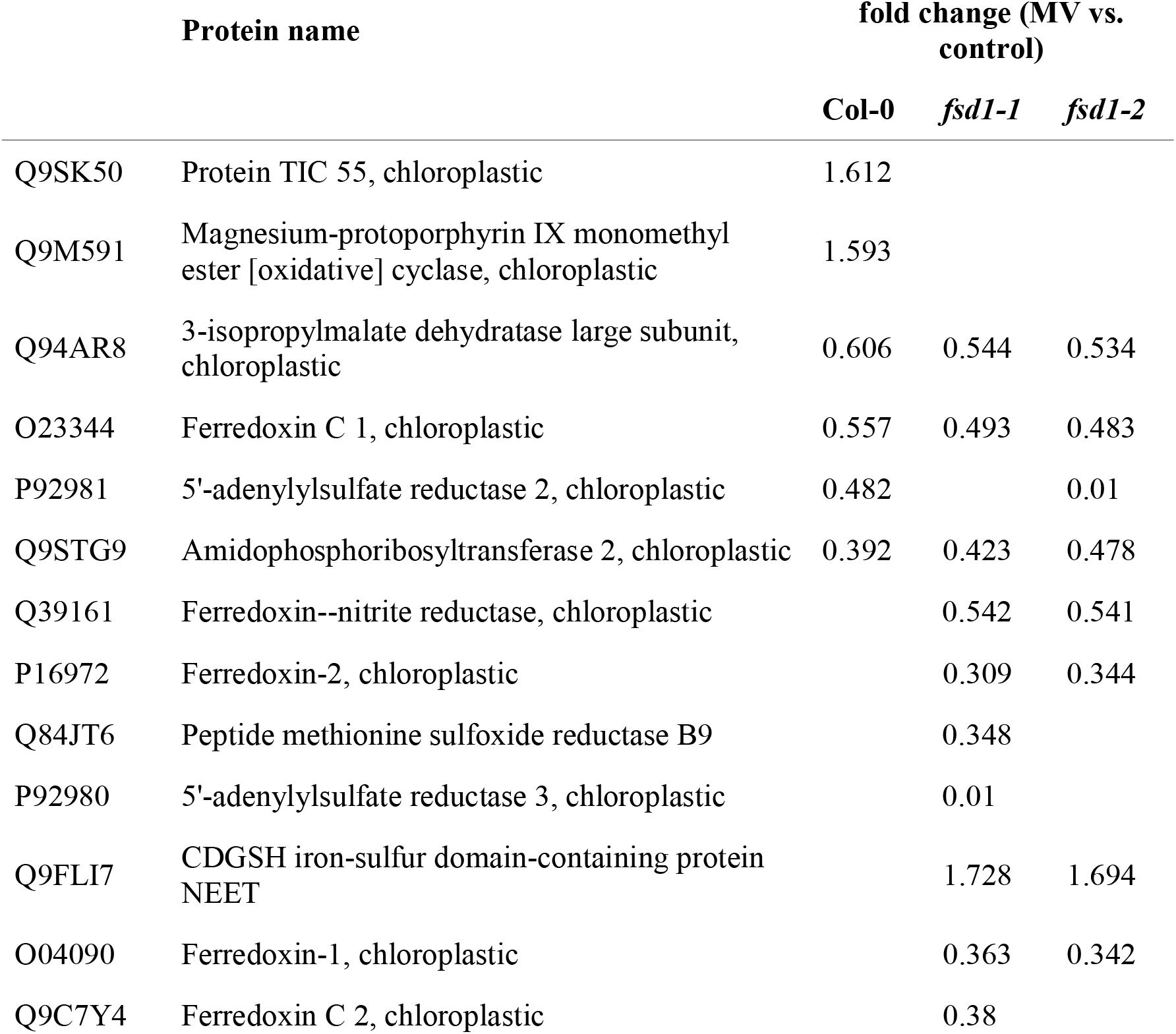
Proteins predicted to possess Fe-S clusters in the differential proteomes of Col-0, *fsd1-1* and *fsd1-2* mutants subjected to methyl viologen (MV). Note that the empty cell means that the protein was not found in the differential proteome of a respective line.

To determine possible functional links among the proteins, we constructed a protein interaction network from the differential proteomes of WT and mutant proteins (Figures S10-12). The MV-responsive differential proteome of WT contains an abundant cluster comprising proteins involved in chloroplast development, including proteins that regulate chloroplast protein import and D1 protein turnover (prevalently lowered abundance). This cluster is connected to photosynthetic proteins and magnesium-protoporphyrin IX monomethyl ester (oxidative) cyclase (increased abundance), which is involved in chlorophyll biosynthesis (Figure S10). The differential proteome of the *fsd1-1* mutant forms an abundant cluster that encompasses photosynthetic proteins together with 5’-adenylylsulfate reductase 3, which is important for cysteine biosynthesis, and pterin-4-alpha-carbinolamine dehydratase (both decreased in abundance), which regulates ribulose 1,5-bisphosphate carboxylase/oxygenase (RuBisCO) assembly (Figure S11). These results indicate a possible alteration in RuBisCO stability in the mutants. A very similar cluster was also identified in the *fsd1-2* differential proteome (Figure S12). All three differential proteomes formed a cluster of ribosomal proteins, which are in the WT interconnected with glutathione transferase F6. In mutants, we encountered higher rates of proteins regulating chloroplastic protein translation, which are prevalently increased (Figures S11, S12).

Both WT and mutant differential proteomes formed clusters comprising proteins involved in glucosinolate biosynthesis. However, unlike WT (Figure S10), a small cluster of fatty acid biosynthesis proteins occurred in *fsd1-1* (Figure S11). Fatty acid biosynthetic genes were found also in the *fsd1-2* proteome (Table S1), nevertheless, they have not been identified by the STRING analysis. These findings showed that the differential regulation of photosynthetic proteins is associated with D1 protein turnover and chloroplast development in WT, and downregulation of RuBisCO assembly, acceleration of translation, and fatty acid biosynthesis in mutants.

Furthermore, we noticed that MV also affected the iron storage protein ferritin 3 in the WT (Table S1). Proteomic data indicated that ferritin levels were altered more strongly in the mutants because they had lower levels of ferritin 1 in addition to ferritin 3. Immunoblotting analysis using anti-ferritin primary antibody demonstrated that MV treatment caused a more intensive decrease in the abundance of ferritins in the mutants than in WT (Figures 7 C, D).

## Discussion

### The ROS-protective function of FSD1 is Cu^2+^ dependent

The current knowledge about the antioxidative roles of FSD1 is controversial [24,28,30,51]. Our analyses indicate that FSD1 plays a protective role in a Cu^2+^-dependent manner. The expression and abundance of FSD1 strictly depend on Cu^2+^ and reach its maximum under Cu^2+^ deficiency [27]. This dependence is linked to the transcriptional control mechanism of *FSD1* expression. A conserved *cis-*element, present in the promoter sequence of *FSD1*, binds the transcription factor SPL7, which activates *FSD1* expression [35]. At the same time, SPL7 suppresses the expression of *CSD1* and *CSD2* by upregulating miRNA398, which negatively regulates these two CSD isoforms [33]. Our results explain the recently shown MV hypersensitivity of *fsd1* mutants grown on half-strength MS media [27], which contains low Cu^2+^ concentrations (0.05 μM) and thus the ratio of FSD1 to CSD activity is high. Most likely, the regulation of O_2_^•–^ levels generated in chloroplasts is altered by the CSD2 isoform under high Cu^2+^ levels. Therefore, we conjectured that the O_2_^•–^ dismutation in chloroplasts is ensured by the strict suborganellar distribution of CSD2 and FSD isoforms and an intricate regulatory mechanism evolved for environments with different Cu^2+^ concentrations.

Differences in the O_2_^•–^ accumulation rate in the leaves of the *fsd1* mutants compared to the WT after MV treatment underline the role of FSD1 in the defense against oxidative stress at low Cu^2+^ concentrations. A more pronounced decrease in O_2_^•–^ production rate after MV treatment in the *fsd1* mutants and the GFP-FSD1 line likely resulted from their reduced formation due to inhibited photosynthesis. Indeed, PAM analysis showed a greater reduction in the quantum efficiency of PSII in the *fsd1* mutants and the GFP-FSD1 line in response to MV, compared to the WT and FSD1-GFP lines. Interestingly, according to the proteomic data, proteins that constitute the LHC of PSII are sensitive to MV in the *fsd1* mutants but not in the WT, suggesting that FSD1 is important for the elimination of ROS, which is detrimental to PSII.

Reportedly, MV induces changes in the structure and turnover of the D1 protein in PSII [14,15]. Therefore, ROS, whose early massive production is catalyzed by MV, are probably responsible for reducing the photochemical efficiency of PSII, which might contribute to the lower O_2_^•–^ production rate after 4 h of MV exposure. Based on this assumption, it can be inferred that *fsd1* mutants and GFP-FSD1 plants lack the protective role of plastidic FSD1 in the initial stages of MV treatment. This is supported by the increased FSD1 activity shortly after MV application in WT plants. Additionally, FSD1 activity did not increase in the line expressing only cytoplasmic and nuclear FSD1, highlighting the role of plastidial FSD1 in the plant oxidative stress response.

Concomitantly with low O_2_^•–^ levels, we observed an overaccumulation of H_2_O_2_, which most likely originates from processes independent of photosynthetic O_2_^•–^ production, such as the oxidation of fatty acids and the activity of oxidases [52]. The elevated H_2_O_2_ was accompanied by the inhibition of APX activity and cAPX abundance in the *fsd1* mutants and GFP-FSD line. The downregulation of APX is most likely a result of highly oxidative conditions after 4 h of MV treatment, as determined by the protein carbonylation assay. In agreement, a previous study showed that MV caused significant oxidative damage to the proteins in *A. thaliana*, which led to the induction of autophagy and thus to the degradation of these damaged proteins [53]. Taken together, we propose that FSD1 prevents H_2_O_2_ accumulation during oxidative stress under limited Cu^2+^ concentrations in plants.

### FSD1 deficiency increases the sensitivity of Fds to MV

Oxidative modifications of proteins have been efficiently monitored using redox proteomics approaches [54,55]. For example, MV causes the oxidation of a wide range of proteins, including RuBisCO, PSII oxygen*-*evolving complexes 1 and 23K, PSII oxygen*-* evolving enhancers PSBQ1 and 2, and glutathione S-transferases in *Arabidopsis* [56]. ROS overaccumulation leads to destructive effects on proteins, affecting turnover [57]. Quantitative differential shotgun proteomics efficiently reflects changes in protein abundance [58,59]. Accordingly, MV in our experimental system led to remarkable changes in *Arabidopsis* WT and *fsd1* proteomes.

FSD1 deficiency deregulates Fe-S cluster*-*binding proteins and proteins involved in Fe-S cluster biogenesis during exposure of plants to MV. Proteins binding to the Fe-S cluster are targeted by O_2_^•–^ and H_2_O_2_ with differing velocities. The oxidation of Fe-S clusters occurs more slowly by H_2_O_2_ than by O_2_^•–^ [60]. Thus, O_2_^•–^ efficiently inhibits the enzymatic activity of Fe-S containing dehydratases, aconitase, transketolase, and other enzymes such as catalase and glutathione peroxidase [61]. The sensitivity of Fe-S cluster binding proteins depends on the stoichiometry of the Fe-S cluster and its exposure to the environment [62]. These findings can explain that some Fe-S cluster-containing proteins showed an increase in abundance in response to MV in our results. The breakdown of the Fe-S cluster elevates the levels of free iron and intensifies the generation of highly reactive ^•^OH by the Fenton reaction [60]. Therefore, the elevated downregulation of Fe-S cluster proteins in the mutants indicates a protective role of FSD1 in Fe-S binding protein stability during oxidative stress.

The most pronounced differences between the mutants and WT were found in the Fd isoforms. Fd is a small 2Fe-2S cluster-containing protein, a stromal electron acceptor in PSI [63]. Fds have a substantial impact on redox regulation and antioxidant defense [63]. Under oxidative stress, the Fe-S cluster of Fd undergoes disassembly [64] and Fd is downregulated in response to environmental cues [65]. Its oxidation may control the distribution of electrons in either linear or cyclic electron flow [63]. Fds 1 and 2 are designated as two chloroplastic isoforms in *Arabidopsis*, out of which ferredoxin 2 constitutes more than 90% of the total leaf ferredoxin complement. Fd1 is believed to be important for cyclic electron flow [66]. Our results showed that FSD1 deficiency sensitized multiple Fd isoforms to MV, suggesting that FSD1 is linked to the abundance of Fds and might protect them from oxidative damage.

Ferritins, the iron storage proteins, represent another known O_2_^•–^ target. They are important for oxidative stress tolerance, and their downregulation elevates ROS levels [67]. O_2_^•–^ reductively mobilizes free iron from ferritins, potentiating the oxidative damage caused by ^•^OH generated by the Fenton reaction [68]. The increased downregulation of ferritin in the mutants compared to WT further supports the role of FSD1 in antioxidative defense.

### FSD1 likely contributes to stress signaling

GO analysis showed that MV treatment caused an altered abundance of proteins involved in pathogen defense in the WT. Among others, it downregulates the cysteine- and histidine-rich domain-containing protein RAR1, which contributes to the plant immunity by supporting the stability of R proteins [69]. Moreover, it also reduces the abundance of RPM1-interacting protein 4 (RIN4), which is targeted by multiple bacterial effectors followed by post-translational modifications, culminating in the suppression of PAMP-triggered immunity [70]. Efficient photosynthesis is important for plant defense against pathogens [71]. It has been shown that chloroplastic H_2_O_2_ induces the expression of genes involved in responses to wounding and pathogen attack [72] and may serve as a messenger in plastid-nucleus retrograde signaling [73]. Alterations in biotic stress-responsive proteins were also reported in a previous transcriptomic study [74]. O_2_^•–^ generated by MV has a broader role during signaling because it activates a wide range of stress-responsive genes [75]. Our data, showing reduced differential regulation of proteins involved in signaling in *fsd1* mutants, suggest the role of FSD1 in the regulation of signaling roles of O_2_^•–^, particularly toward the abundance of biotic stress-responsive proteins. Furthermore, for yeast SOD1, the role of nuclear FSD1 as a putative co-transcription factor has been shown to regulate the signaling roles of O_2_^•–^ [76]. However, further studies are required to ascertain its roles in *Arabidopsis*.

## Conclusions

Our results demonstrate that FSD1 is an important component of antioxidant machinery, protecting plants against the detrimental effects of O_2_^•–^ under low Cu^2+^ concentrations in the growth substrate. It contributes to sustained levels of Fe-S cluster-binding proteins, particularly Fd isoforms, under increased chloroplast ROS production.

## Acknowledgements

This research was funded by Grant No. 19-00598S from the Czech Science Foundation GAC□R and by the ERDF project “Plants as a tool for sustainable global development” (No.CZ.02.1.01/0.0/0.0/16_019/0000827). Mass spectrometry analyses were performed at the Turku Proteomics Facility supported by Biocenter Finland. We thank Constance Journet for the help in chlorophyll fluorescence imaging.

## Author Contributions

PM, PD, YK and AS conducted the experiments and drafted the manuscript with input from all co-authors. TT, JŠ and JK revised and edited the manuscript. TT conceived and supervised the project.

## Data availability

The mass spectrometry proteomics data have been deposited to the ProteomeXchange Consortium via the PRIDE [77] partner repository with the dataset identifier PXD028328.

