## Supplemental Figures for "*Arabidopsis* iron superoxide dismutase 1 protects against methyl viologen-induced oxidative stress in a copper-dependent manner"

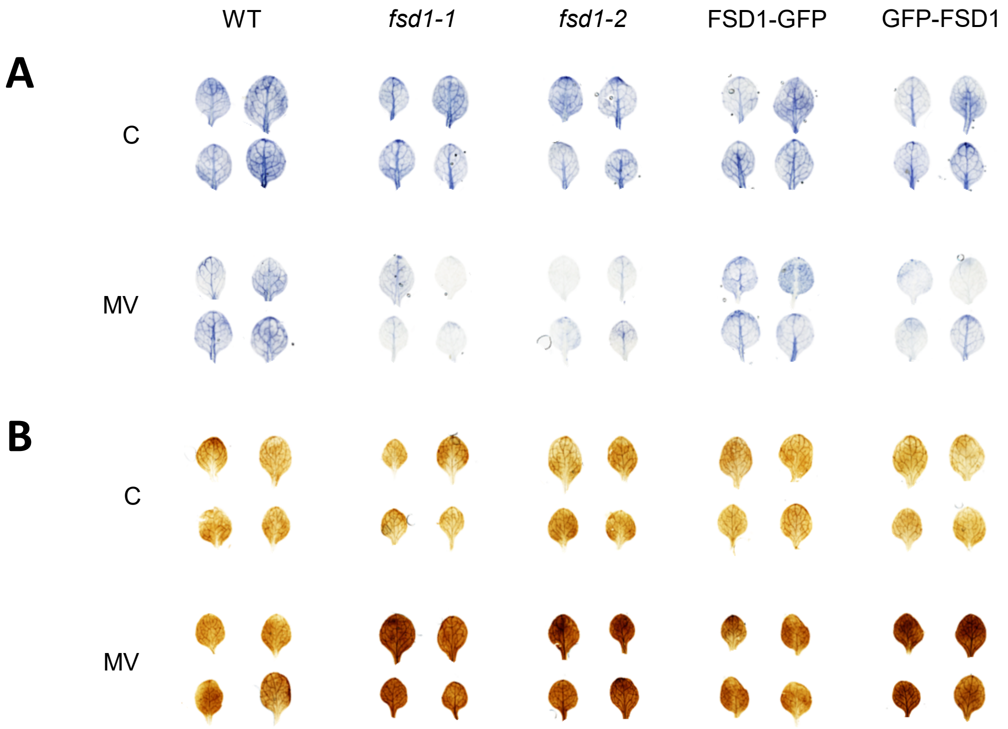


Figure S1. Generation of superoxide ($O_2^{\bullet-}$) and hydrogen peroxide ($H_2O_2$) in whole leaf rosettes of 10-day-old *Arabidopsis* plants of wild type WT, *fsd1* mutants, FSD1-GFP and GFP-FSD1 lines. $O_2^{\bullet-}$ production was visualized as dark blue coloration by nitroblue tetrazolium (NBT) staining (A), and $H_2O_2$ production was visualized as dark brown coloration by 3,3′-diaminobenzidine (DAB) staining (B).

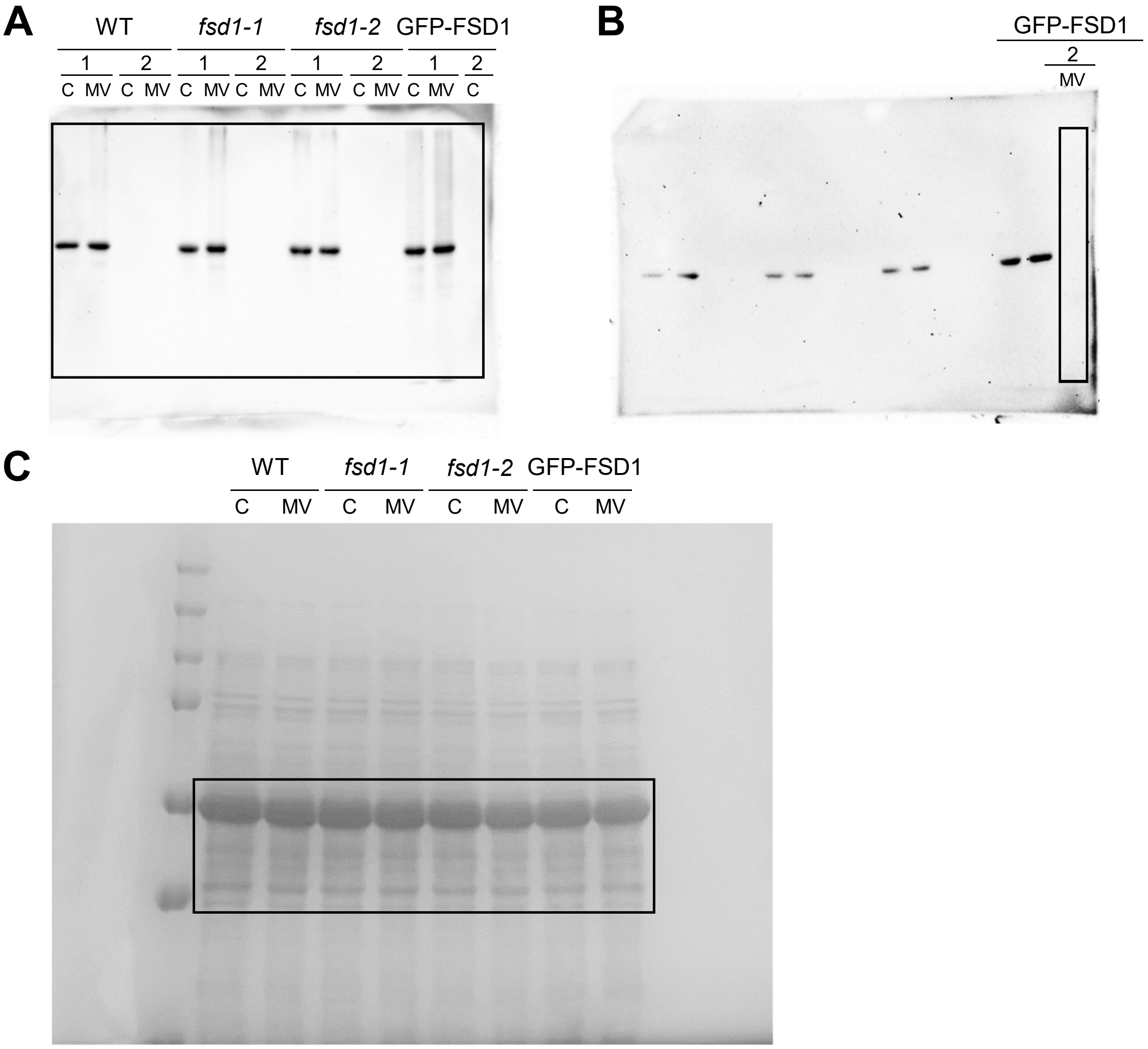


Figure S2. Full scan of the entire original blots and Comassie Briliant Blue G-250 stained gel presented in Figure 3A. (A, B) Entire membranes with chemiluminiscent signal observed after probing with anti-DNP antibody included in OxyBlot$^{TM}$ kit. Each blot contains protein extracts from mock (lane C) and 1 μM MV treated (lane MV) plants. Each sample was treated either with DNPH oxidation reagent (1) or a control reagent (2) included in the OxyBlot kit. (B) The MV treated sample of GFP-FSD1 derivatized with control reagent was run on a separate gel (B) followed by the same procedure applied for A. Samples loaded on lanes which are not annotated are not relevant to this study. The highlighted regions show the presented sections in Figure 3A (C) Entire Comassie Briliant Blue G-250 stained gel. The highlighted region shows the presented section in Figure 3C.

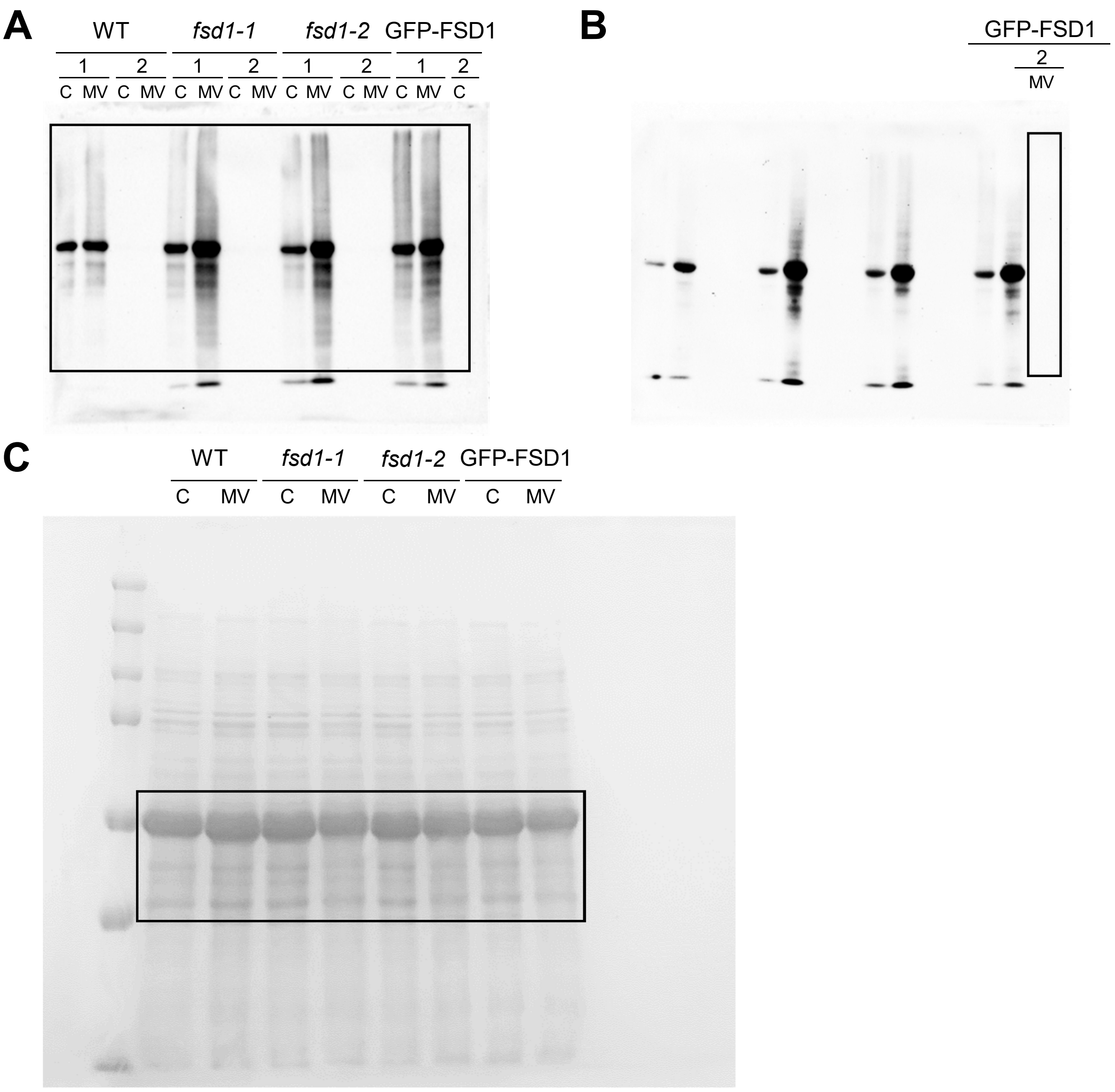


Figure S3. Full scan of the entire original oxyblots and Comassie Briliant Blue G-250 stained gel presented in Figure 3B. (A, B) Entire membranes with chemiluminiscent signal observed after probing with anti-DNP antibody included in OxyBlot$^{TM}$ kit. Each blot contains protein extracts from mock (lane C) and 1 $\mu$M MV treated (lane MV) plants. Each sample was treated either with DNPH oxidation reagent (1) or a control reagent (2) included in the OxyBlot kit. (B) The MV treated sample of GFP-FSD1 derivatized with control reagent was run on a separate gel (B) followed by the same procedure applied for membrane A. Samples loaded on lanes which are not annotated are not relevant to this study. The highlighted regions show the presented sections in Figure 3B (C) Entire Comassie Briliant Blue G-250 stained gel. The highlighted region shows the presented section in Figure 3D.

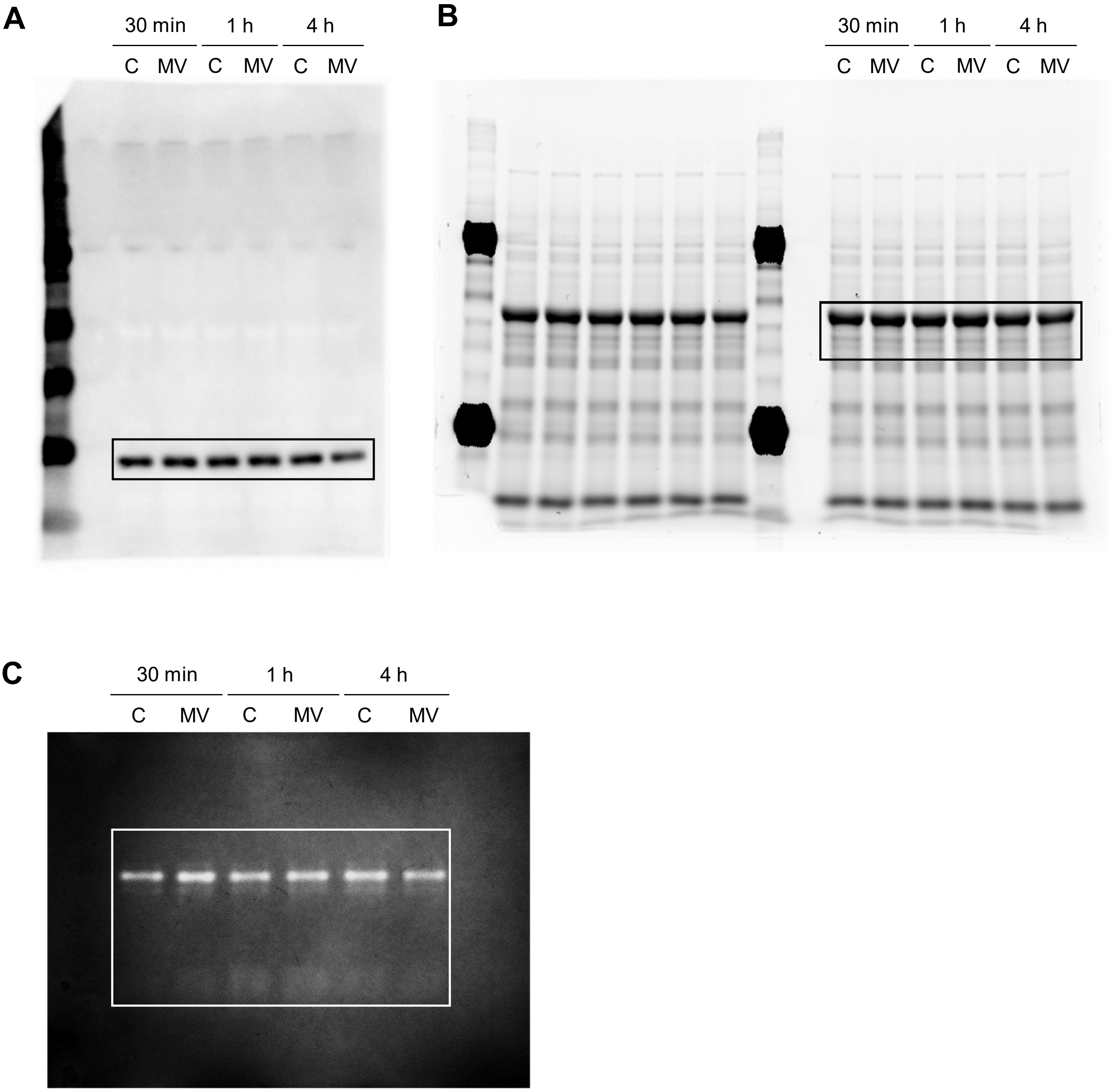


Figure S4. Full scan of the entire original immunoblot and gel stained for specific activity of superoxide dismutases presented in Figure 4. (A) Entire membrane with chemiluminiscent signal observed after probing with anti-FSD1 antibody. (B) Full image of the Stain-free gel with separated proteins representing equal amount of proteins loaded on gel prior to transfer to PVDF membrane. The highlighted regions show the sections presented in Figure 4A. Samples loaded on lanes which are not annotated are not relevant to this study. (C) Full scan of the entire original gel stained for specific activity of superoxide dismutases. The highlighted region shows the presented section in Figure 4C. The membrane (A) and gels (B, C) contain protein extracts from mock control conditions (lane C) and 1 μM MV treatment (lane MV) treated for 30 min, 1 h and 4 h.

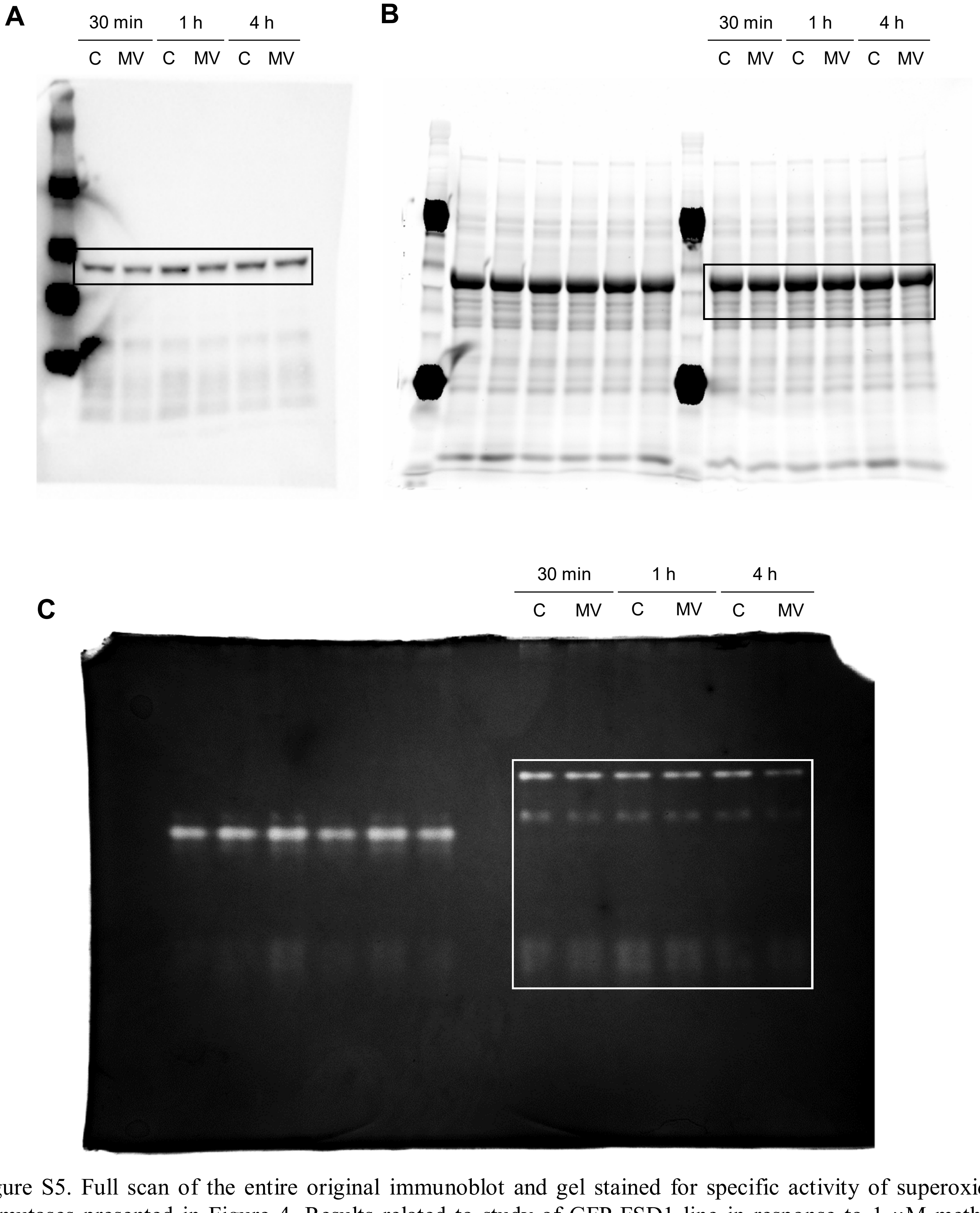


Figure S5. Full scan of the entire original immunoblot and gel stained for specific activity of superoxide dismutases presented in Figure 4. Results related to study of GFP-FSD1 line in response to 1 μM methyl viologen (MV) are shown. (A) Entire membrane with chemiluminiscent signal observed after probing with anti-FSD1 antibody. (B) Full image of the Stain-free gel with separated proteins representing equal amount of proteins loaded on gel prior to transfer to PVDF membrane. The highlighted regions show the sections presented in Figure 4E. Samples loaded on lanes which are not annotated are not relevant to this study. (C) Full scan of the entire original gel stained for specific activity of superoxide dismutases. The highlighted region shows the presented section in Figure 4G. The membrane (A) and gels (B, C) contain protein extracts from mock control conditions (lane C) and 1 μM MV treatment (lane MV) treated for 30 min, 1 h and 4 h.

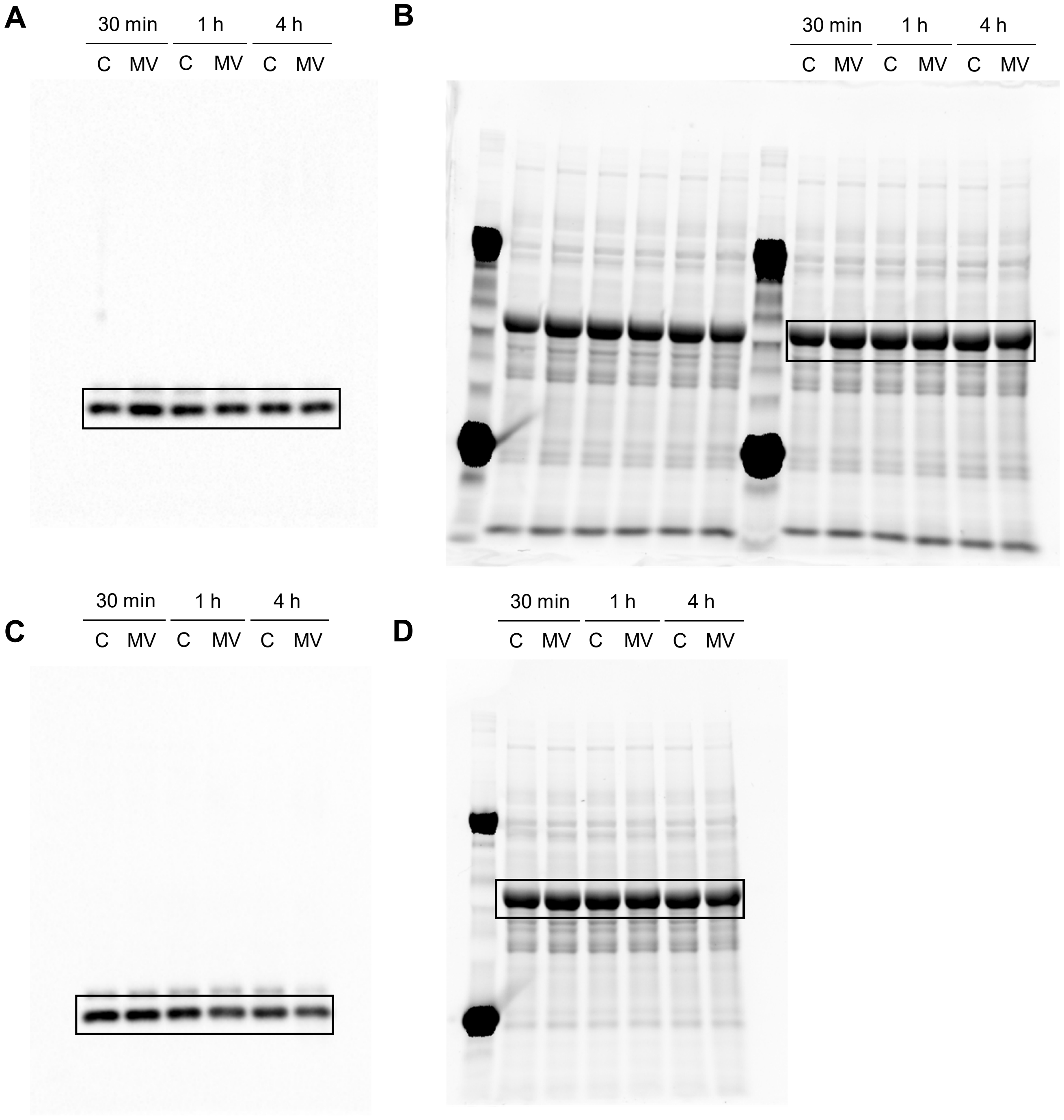


Figure S6. Full scan of the entire original immunoblots presented in Figure 5. Results related to wild type (WT) (A, B) and *fsd1-1* mutant (C, D) in response to 1 µM methyl viologen (MV) are shown. (A, C) Entire membrane with chemiluminiscent signal observed after probing with anti-APX antibody. (B, D) Full image of the Stain-free gel with separated proteins representing equal amount of proteins loaded on gel prior to transfer to PVDF membrane. The highlighted regions show the sections presented in Figure 5B (A, B) and 5C (C, D). Samples loaded on lanes which are not annotated are not relevant to this study. Membranes (A, C) and gels (B, D) contain protein extracts from mock control conditions (lane C) and 1 µM MV treatment (lane MV) treated for 30 min, 1 h and 4 h.

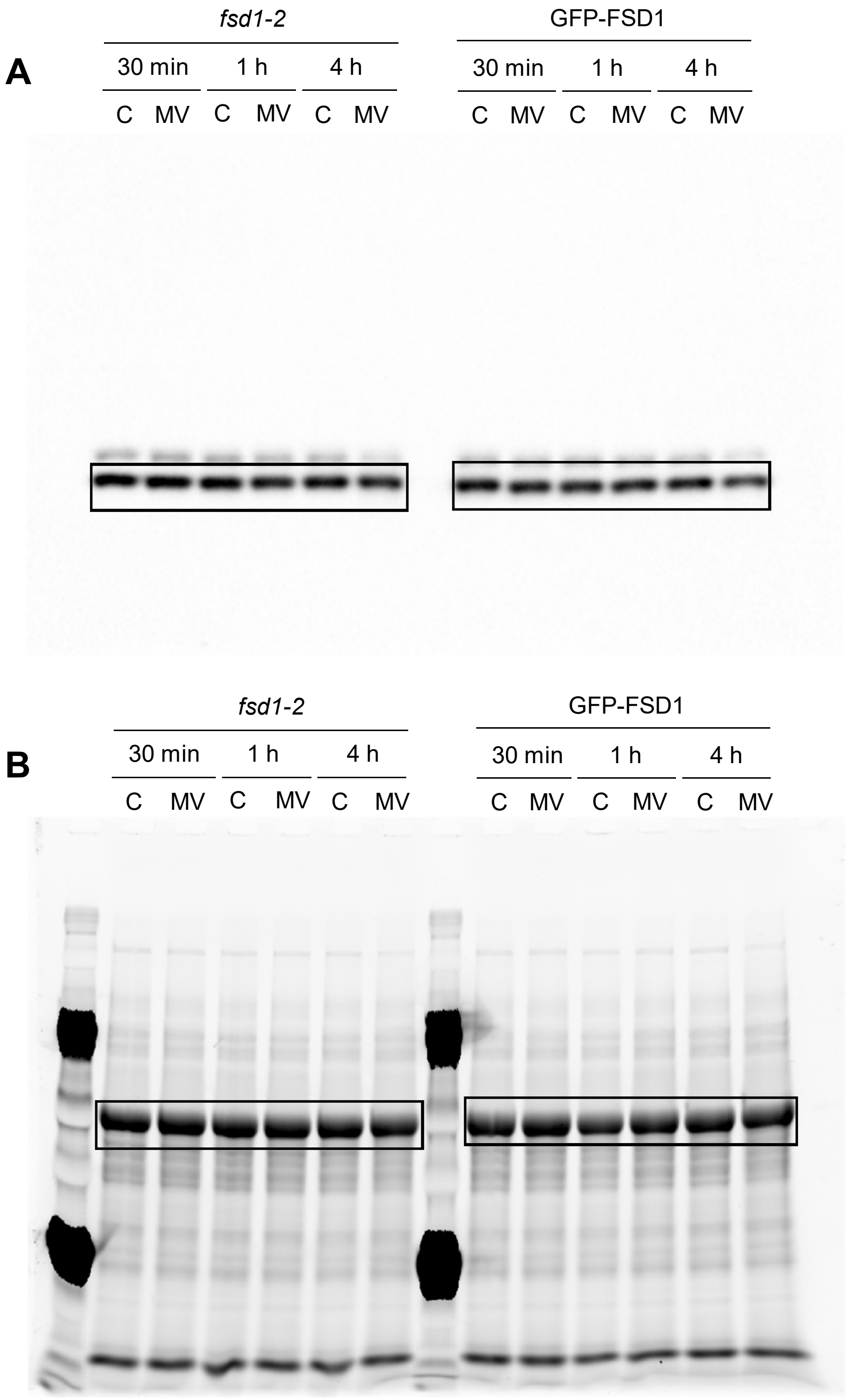


Figure S7. Full scan of the entire original immunoblots presented in Figure 5. Results related to study of *fsd1-2* mutant and GFP-FSD1 line in response to 1 µM methyl viologen (MV) are shown. (A) Entire membrane with chemiluminiscent signal observed after probing with anti-APX antibody. (B) Full image of the Stain-free gel with separated proteins representing equal amount of proteins loaded on gel prior to transfer to PVDF membrane. The highlighted regions show the sections presented in Figure 5D and 5E. Membrane and gel contain protein extracts from mock control conditions (lane C) and 1 µM MV treatment (lane MV) treated for 30 min, 1 h and 4 h.

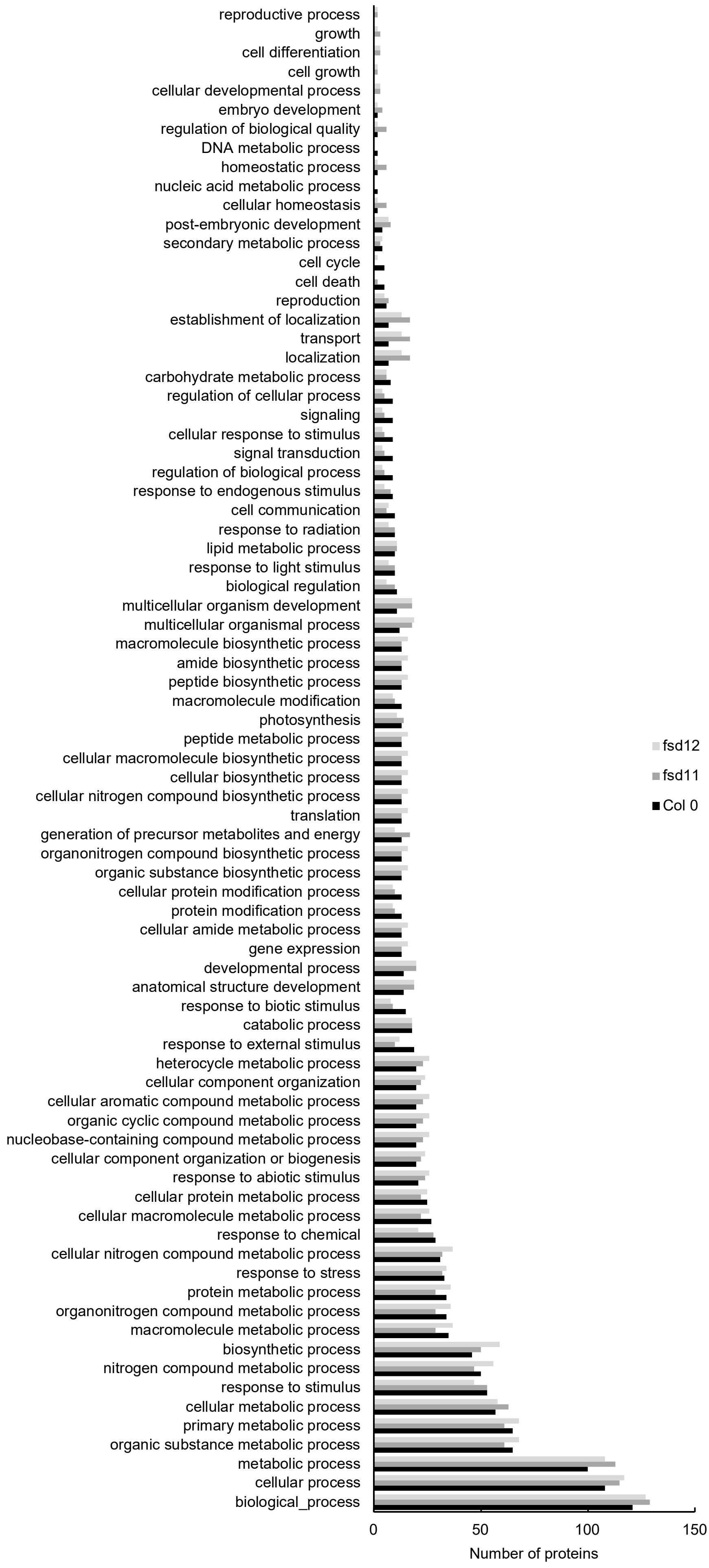


Figure S8 showing the abundance of gene ontology annotations found in the differential proteomes of the examined lines according to biological process .

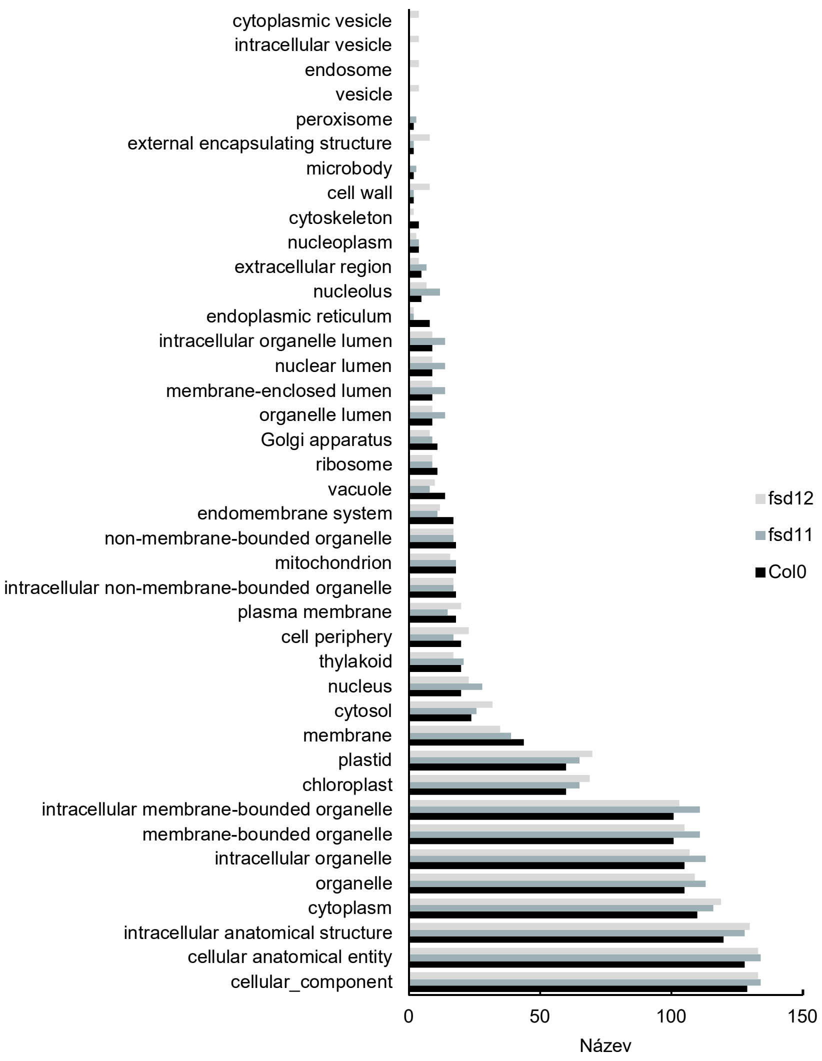


Figure S9. Graph showing the abundance of gene ontology annotations found in the differential proteomes of the examined lines according to cell compartment.

| Chloroplast protein import, quality control and chloroplast development | Ratio |
|---|---|
| Ferredoxin C 1 | 0.56 |
| Probable zinc metalloprotease EGY2 | 2.28 |
| Thylakoid lumenal 15.0 kDa protein 2 | 1.59 |
| Threonine--tRNA ligase 2 | 0.52 |
| Protein BUNDLE SHEATH DEFECTIVE 2 | 0.54 |
| ATP-dependent zinc metalloprotease FTSH 1 | 0.48 |
| ATP-dependent zinc metalloprotease FTSH 2 | 0.57 |
| Protein TIC 55 | 1.61 |

| Small Heat shock proteins | Ratio |
|---|---|
| 23.5 kDa heat shock protein, mitochondrial | 0.01 |
| 17.6 kDa class I heat shock protein 2 | 0.01 |

| ER protein quality control | Ratio |
|---|---|
| Heat shock 70 kDa protein 17 | 0.55 |
| Stromal cell-derived factor 2-like protein | 0.34 |

| Photosynthetic proteins | Ratio |
|---|---|
| Plastocyanin major isoform | 1.62 |
| Photosystem I chlorophyll a/b-binding protein 3-1 | 1.61 |
| Magnesium-protoporphyrin IX monomethyl ester [oxidative] cyclase | 1.59 |
| Chlorophyll a-b binding protein 2 | 1.59 |
| Chlorophyll a-b binding protein 2.1 | 1.62 |
| Photosystem II reaction center protein H | 1.65 |

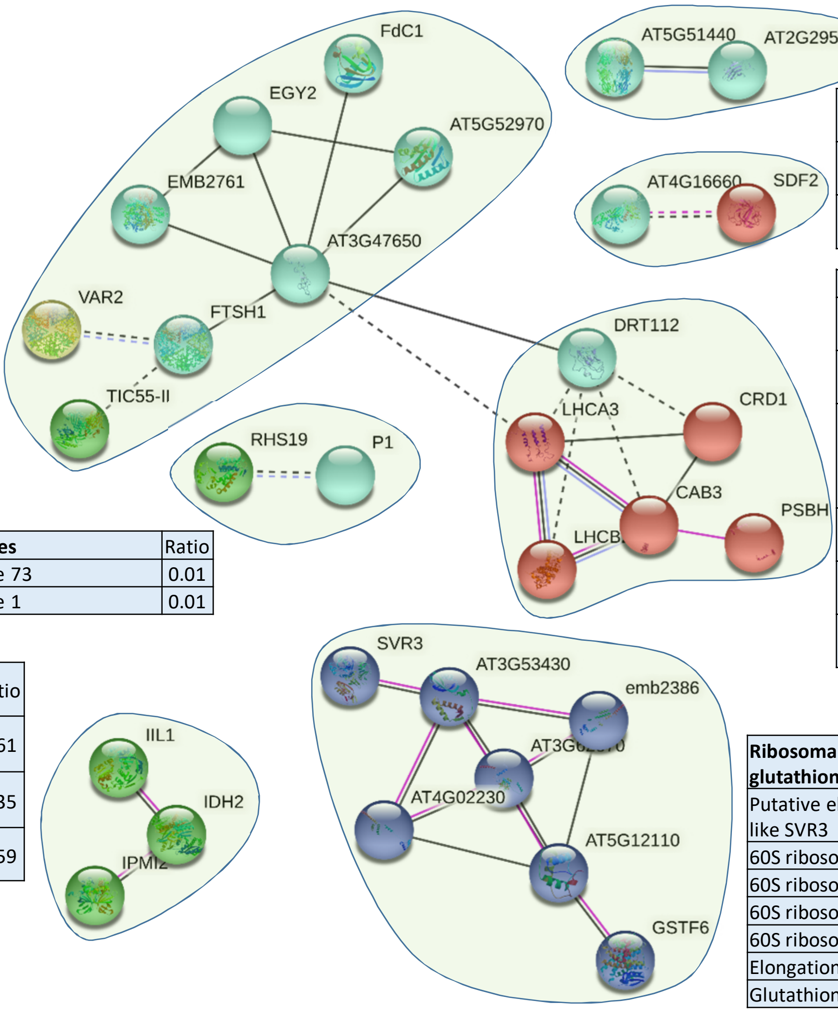


| Peroxidases | Ratio |
|---|---|
| Peroxidase 73 | 0.01 |
| Peroxidase 1 | 0.01 |

| Metabolism , glucosinolate biosynthesis, | Ratio |
|---|---|
| 3-isopropylmalate dehydratase large subunit | 0.61 |
| Isocitrate dehydrogenase [NAD] regulatory subunit 2 | 0.35 |
| 3-isopropylmalate dehydratase small subunit 2 | 0.59 |

| Ribosomal proteins and glutathione transferase | Ratio |
|---|---|
| Putative elongation factor TypA-like SVR3 | 0.61 |
| 60S ribosomal protein L12-2 | 0.15 |
| 60S ribosomal protein L19-1 | 0.51 |
| 60S ribosomal protein L7a-2 | 0.61 |
| 60S ribosomal protein L19-3 | 0.55 |
| Elongation factor 1-beta 1 | 0.56 |
| Glutathione S-transferase F6 | 0.63 |

Figure S10. Schematic presentation of protein interaction networks in differential proteome of MV-treated wild type as generated by STRING web-based application.

| Photosynthesis | Ratio |
|---|---|
| Ferredoxin C 1 | 0.49 |
| Probable pterin-4-alpha-carbinolamine dehydratase | 0.61 |
| 5'-adenylylsulfate reductase 3 | 0.01 |
| Ferredoxin-1 | 0.36 |
| Ferredoxin-2 | 0.31 |
| Ferredoxin--nitrite reductase | 0.54 |
| Glucose-6-phosphate 1-dehydrogenase 3 | 100.00 |
| Photosystem I reaction center subunit VI-1 | 31.10 |
| 50S ribosomal protein 5 | 0.62 |
| 31 kDa ribonucleoprotein | 1.69 |

| Fatty acid biosynthesis | Ratio |
|---|---|
| 3-oxoacyl-[acyl-carrier-protein] synthase II | 0.58 |
| Acyl carrier protein 2 | 1.68 |

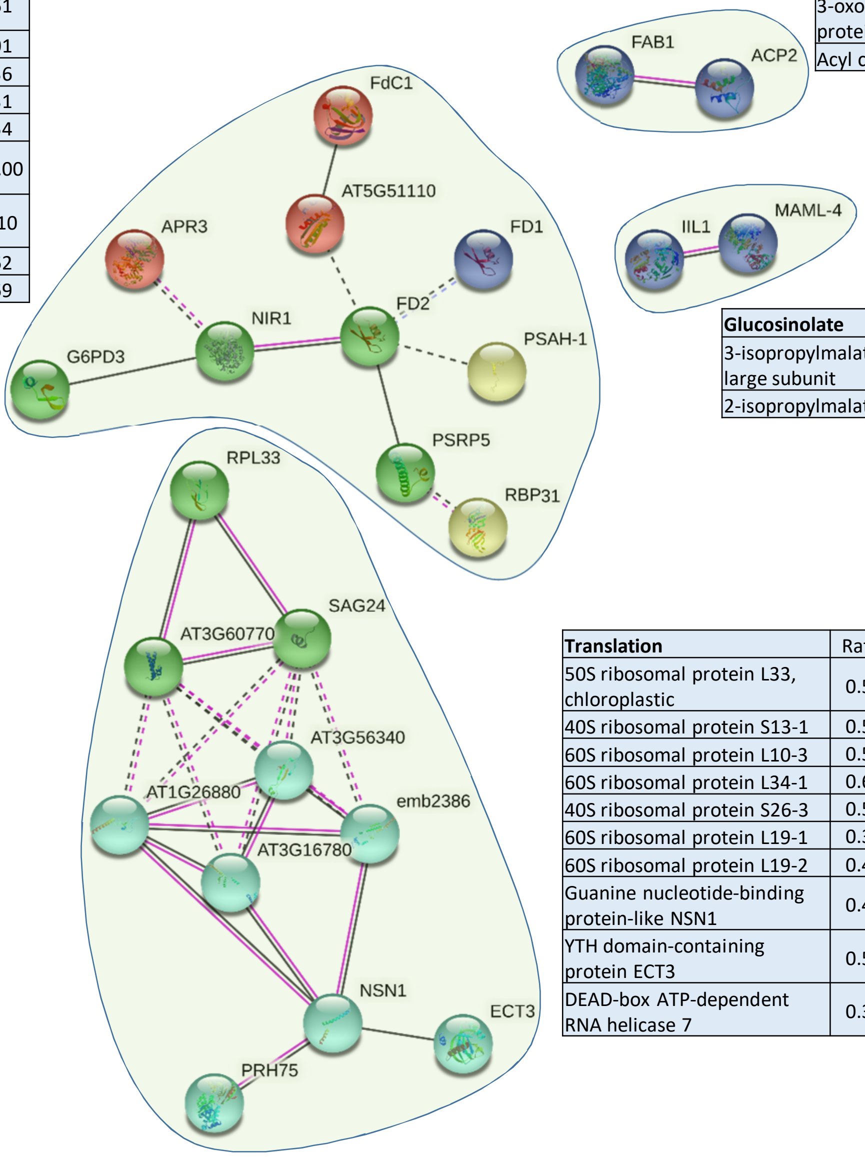


| Glucosinolate | Ratio |
|---|---|
| 3-isopropylmalate dehydratase large subunit | 0.54 |
| 2-isopropylmalate synthase 1 | 0.63 |

| Translation | Ratio |
|---|---|
| 50S ribosomal protein L33, chloroplastic | 0.52 |
| 40S ribosomal protein S13-1 | 0.50 |
| 60S ribosomal protein L10-3 | 0.52 |
| 60S ribosomal protein L34-1 | 0.63 |
| 40S ribosomal protein S26-3 | 0.59 |
| 60S ribosomal protein L19-1 | 0.38 |
| 60S ribosomal protein L19-2 | 0.45 |
| Guanine nucleotide-binding protein-like NSN1 | 0.47 |
| YTH domain-containing protein ECT3 | 0.50 |
| DEAD-box ATP-dependent RNA helicase 7 | 0.38 |

Figure S11. Schematic presentation of protein interaction networks in differential proteome of MV-treated *fsd1-1* as generated by STRING web-based application.

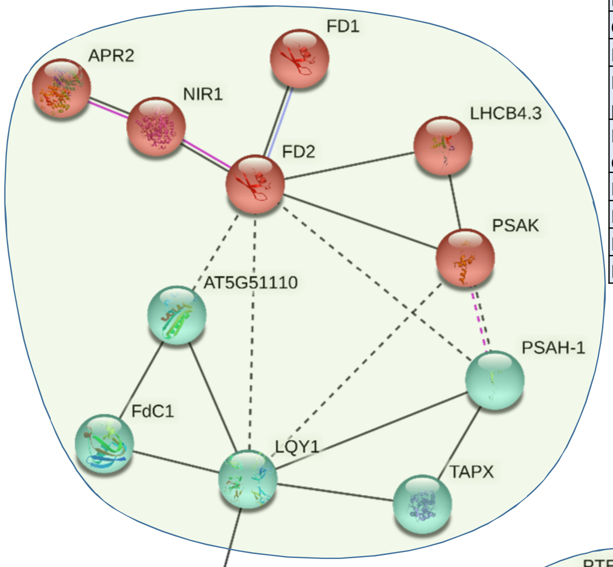


| **Photosynthetic proteins** | Ratio |
|---|---|
| Ferredoxin-1 | 0.34 |
| 5'-adenylylsulfate reductase 2 | 0.01 |
| Ferredoxin--nitrite reductase | 0.54 |
| Chlorophyll a-b binding protein CP29.3 | 0.52 |
| Ferredoxin-2, chloroplastic | 0.34 |
| Photosystem I reaction center subunit psaK | 1.62 |
| Probable pterin-4-alpha-carbinolamine dehydratase | 0.59 |
| Photosystem I reaction center subunit VI-1 | 1.48 |
| Ferredoxin C 1, chloroplastic | 0.48 |
| Protein disulfide-isomerase LQY1 | 0.51 |
| L-ascorbate peroxidase T | 0.55 |

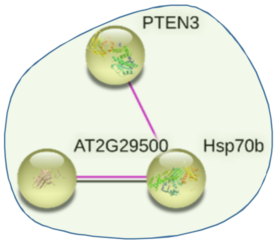


| **Lipid signalling and HSPs** | Ratio |
|---|---|
| Phosphatidylinositol 3,4,5-trisphosphate 3-phosphatase and protein-tyrosine-phosphatase PTEN2B | 2.58 |
| 17.6 kDa class I heat shock protein 2 | 0.01 |
| Heat shock 70 kDa protein 5 | 100.00 |

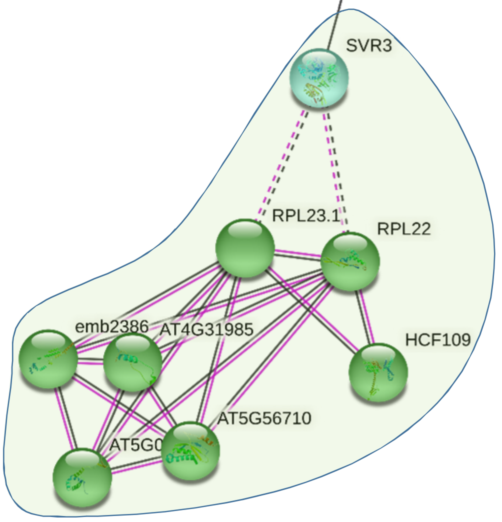


| **Ribosomal proteins** | Ratio |
|---|---|
| Putative elongation factor TypA-like SVR3, chloroplastic | 0.57 |
| 50S ribosomal protein L23, chloroplastic | 2.11 |
| 50S ribosomal protein L22, chloroplastic | 1.57 |
| 60S ribosomal protein L19-1 | 0.39 |
| 60S ribosomal protein L39-1 | 0.56 |
| Peptide chain release factor PrfB1, chloroplastic | 0.50 |
| 60S ribosomal protein L31-3 | 1.56 |
| 40S ribosomal protein S17-4 | 100.00 |

Figure S12. Schematic presentation of protein interaction networks in differential proteome of MV-treated *fsd1-2* as generated by STRING web-based application.

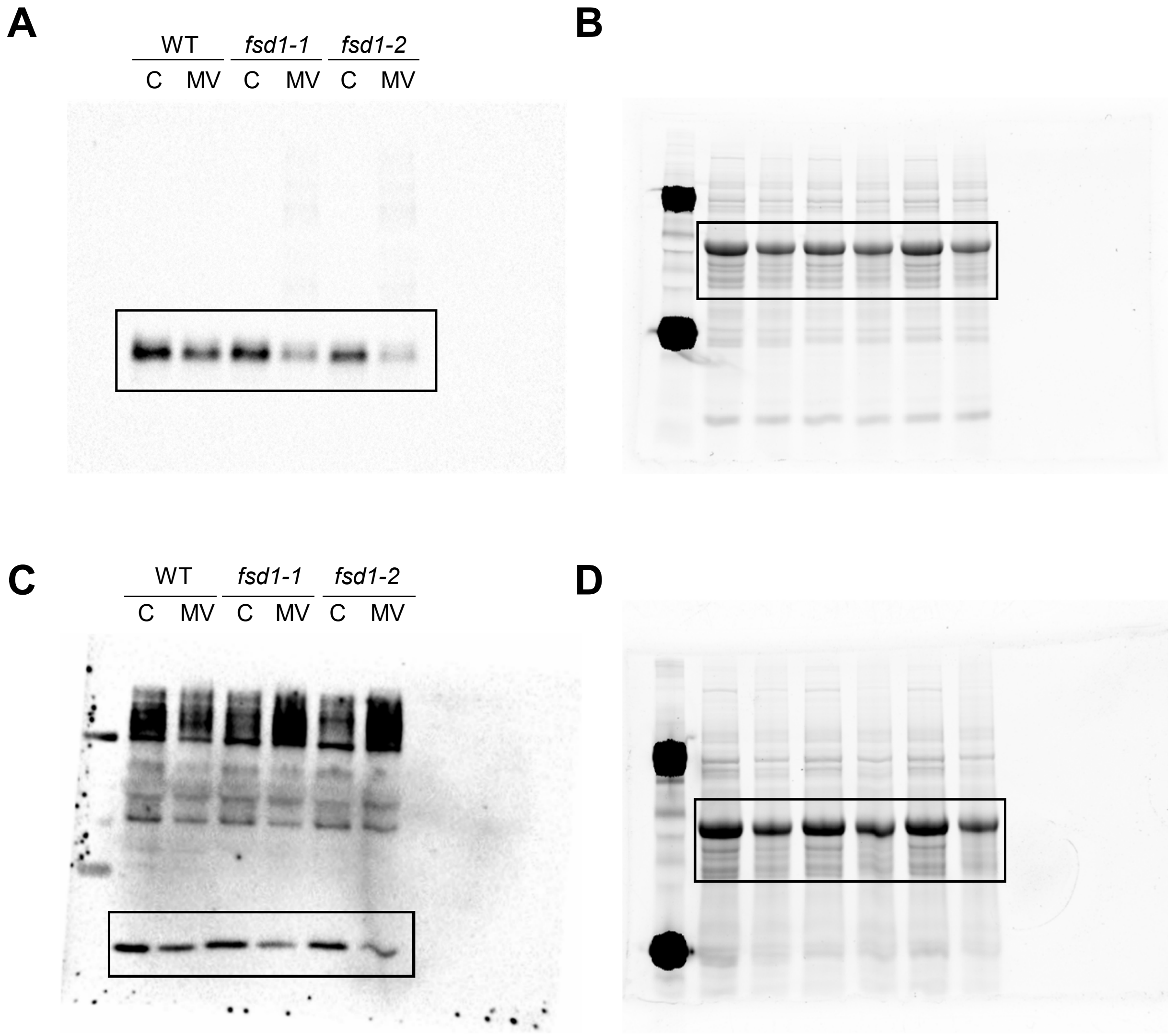


Figure S 13 Full scan of the entire original immunoblots presented in Figure 7. (A) Entire membrane with chemiluminiscent signal observed after probing with anti ferredoxin 2 antibody. The highlighted region shows the section presented in Figure 7A. (B) Respective controls of protein loading using Stain-free gels and the highlighted region shows the section presented in Figure 7 A. (C) Entire membrane with chemiluminiscent signal observed after probing with anti ferritin 3 antibody. The highlighted region shows the section presented in Figure 7C. (D) Respective controls of protein loading using Stain-free gels and the highlighted region shows the section presented in Figure 7C.
